# Rapidly and reproducibly building a comprehensive catalogue of resistance-associated variants for *M. tuberculosis* surveillance

**DOI:** 10.1101/2025.10.02.679941

**Authors:** Dylan Adlard, Kerri M Malone, Jeremy Westhead, Martin Hunt, Hieu Thai, Matthew Colpus, Robert D Turner, Shaheed V Omar, David W Eyre, Nazir Ismail, Timothy M Walker, Timothy EA Peto, Derrick W Crook, Zamin Iqbal, Philip W Fowler

**Author notes:** To whom correspondence should be addressed, @philipwfowler.bsky.social.

## Abstract

**Background:** Catalogues of genetic variants associated with resistance underpin whole-genome sequencing (WGS)- based predictions of drug susceptibility in *Mycobacterium tuberculosis*, and are essential for developing molecular diagnostics and surveillance. The current gold standard catalogues are those released by the WHO but the underlying data are neither fully available nor straightforward to interpret. Open and reproducible methods would help address these problems, extending the important work already done to better prepare for implementation into readily updatable surveillance platforms.

**Methods:** We have developed an automated method, `catomatic`, that uses a binomial test to associate informative isolates with resistance or susceptibility, and built a catalogue (`catomatic-1`) from the same 39,358 samples used to construct the first edition of the WHO catalogue (`WHOv1`). We performed a sensitivity analysis to optimise the statistical and bioinformatic parameters for each drug, and benchmarked `catomatic-1` against `WHOv1` using a hold-out Validation Dataset of 14,380 isolates.

**Findings:** By using simpler statistics, `catomatic-1` algorithmically classified 1,329 genetic variants, ranging from five for linezolid to 440 for pyrazinamide. `WHOv1` included generalisable rules added by a panel of experts, increasing its predictive coverage, but at the cost of automation and deterministic reproducibility. Despite not including such expert rules, `catomatic-1` achieves comparable performance for all drugs, with sensitivities for first-line agents above 88% on the hold-out Validation Dataset. The automated process allowed us to efficiently explore parameter space; for instance, detecting resistance-associated variants with low read support improved the sensitivity for all drugs.

**Interpretation:** Performant resistance catalogues for *M. tuberculosis* can be built automatically and reproducibly using transparent statistical methods. As more data are collected, catalogue content and performance will evolve, highlighting the need for proper versioning, machine/human readability, and open access. This approach demonstrates resistance catalogues used in surveillance (and potentially diagnostics) can be rapidly and reproducibly updated.

**Funding:** The National Institute for Health and Care Research (NIHR), Engineering and Physics Sciences Research Council (EPSRC), the Ellison Institute of Technology (EIT), Oxford Ltd and ORACLE Corporation.

**Research in context:** *Evidence before this study:* We searched PubMed and preprint servers (bioRxiv, medRxiv), and publicly available mutation catalogues for studies linking *Mycobacterium tuberculosis* genomic variants with drug resistance using whole-genome or targeted sequencing and phenotypic drug-susceptibility testing (pDST). Search terms combined “Mycobacterium tuberculosis”, “genome sequencing”, “mutation catalogue”, ”mutation effects”, “drug resistance”, and individual drug names, with no language or date restriction. We included studies providing paired, clinical genomic and pDST or MIC data, excluding purely *in silico* or case-only reports. This work directly builds on methodologies and data published by five prior studies, including our previous work that introduced our method by applying it to a single drug (bedaquiline), and makes primary comparisons with the First (`WHOv1`) and Second (WHOv2) Editions of the WHO Catalogue of mutations in *Mycobacterium tuberculosis*.

*Added value of this study:* Historically, catalogues of resistance mutations in *M. tuberculosis* have been built by combining rules derived from statistical association testing, the literature and the views of experts with the aim of maximising performance. As the sizes of the datasets used to train such catalogues has increased, the relative contributions of each of these sources to performance has shifted. The current gold standard is WHOv2 which applies a conservative statistical association framework to each genetic variant; this approach is well-suited to molecular test design as classifications are unlikely to change with more data but is less appropriate for surveillance of antimicrobial resistance, which requires predictions that directly reflect the underlying data and can be readily updated and systematically parsed. We developed `catomatic`, an open source tool for reproducibly building catalogues of resistance-associated genetic variants. Trained on the same samples used to build `WHOv1` and benchmarked on a hold-out Validation Dataset, `catomatic` achieves comparable definite prediction rates, sensitivity and specificity to `WHOv1` without expert-rule augmentation and despite using simpler statistics. Here we establish it can be used to build a catalogue for multiple antibiotics by introducing per-drug parameter optimisation, thereby demonstrating applicability across diverse resistance mechanisms. It produces machine-readable outputs (CSV/JSON), and we demonstrate that lowering read-support thresholds can improve detection of minor resistance subpopulations.

*Implications of all the available evidence:* Catalogues of resistance-associated variants for *M. tuberculosis* can be rapidly and transparently constructed. Making catalogues available in human/machine-readable formats with uncertainty estimates will improve uptake of WGS for *M. tuberculosis* surveillance and diagnostics; using a reproducible process permits diagnostic test manufacturers, researchers, clinical and public health laboratories to select the level of statistical support necessitated by their specific use-case. Policymakers should balance the benefits of expert rules against loss of automation and end-to-end reproducibility. Future work will expand the size of the datasets used, integrate minimum inhibitory concentration data, and establish consensus workflows for routine, transparent catalogue updates.

## Introduction

Tuberculosis is the poster child for the translation of whole genome sequencing (WGS) into clinical microbiology; genetics is faster, potentially more accurate than phenotypic drug susceptibility testing (pDST), more comprehensive than molecular tests and also yields epidemiological insights. The key to predicting from a *Mycobacterium tuberculosis (Mtb)* genome whether a tuberculosis infection is susceptible or resistant to a range of antibiotics is currently the *catalogue* – a list of known genetic variants and their effects on individual antibiotics ^1^.

The release of the first edition of the catalogue of mutations in *Mycobacterium tuberculosis* complex (MTBC) and their association with resistance by the World Health Organization (WHO) in 2021^2,3^ marked a milestone in the adoption of WGS in tuberculosis diagnostics; we call this catalogue `WHOv1`. A dataset of 41,137 clinical isolates with WGS and pDST data donated by the CRyPTIC and Seq&Treat consortia was analysed ^3^, resulting in 1,486 genetic variants associated with either resistance or susceptibility to one of 13 antituberculars. Following the endorsement of the oral six-month BPaLM regimen for multidrug-resistant TB ^4^, a second edition (WHOv2) was published in late 2023 based on 61,986 clinical isolates ^5^, notably adding resistance-associated variants (RAVs) for bedaquiline (BDQ) and clofazimine (CFZ). Both catalogues were constructed by associating binary phenotypes (resistant or susceptible) to solitary mutations in resistance genes, followed by confidence grading and the addition of rules derived from the scientific literature, previous WHO reports, and pooled evidence, as agreed by an expert panel ^2,5^.

The WHO catalogue’s primary use is to guide the development of molecular tests that detect specific genetic variants. However, catalogues also have an underutilised role in WGS-based surveillance, where they can enable the rapid discovery, classification, and tracking of clinically relevant AMR determinants as new data become available. Despite representing a significant step towards standardising the clinical interpretation of tuberculosis genetics, existing catalogues are not readily applicable to surveillance, owing to several limitations.

Firstly, although expert-derived rules enable predictions for some rare or previous unobserved variants, they do not necessarily reflect the underlying data. They also add complexity to an already resource-intensive and time-consuming process, making routine updates as new data become available a major undertaking. Thirdly, users must grapple with a lack of transparency and reproducibility: although the donated dataset for `WHOv1` is available via the CRyPTIC project ^6^, the `WHOv2` dataset is not public, and, while some code is available ^7^, it is not sufficient to repeat the entire process. Furthermore, since the release of `WHOv2` in November 2023, minor errors in the Excel spreadsheet have been corrected, but these changes are only flagged via the GitHub page ^7^, leading to the risk that people continue to use deprecated versions.

Finally, despite efforts to integrate the WHO catalogues into pipelines and software ^1,8^, their complexity makes them difficult to read and parse ^8^, as reflected in disagreements between validation studies ^9^. This stems from the lack of a single machine-readable artefact, with rules instead distributed between entries in an Excel spreadsheet and natural language descriptions in the accompanying English report.

In this paper, we describe how we have reproducibly built a deterministic catalogue of MTBC genetic variants associated with (or not associated with resistance) to 15 antibiotics using the same dataset that was used to build `WHOv1`. We then demonstrate comparable performance of our catalogue to `WHOv1` using a non-overlapping hold-out subset of 14,380 samples dervied from the latest version (v3.4.0) of the CRyPTIC dataset. This work is enabled by (i) the continued release of versioned data by the CRyPTIC consortium ^10^, (ii) a publicly-available software tool, `catomatic`, that automatically classifies the effect of individual solitary mutations in resistance genes ^11^ resulting in (iii) a catalogue described by a single file that is both human- and computer-parsable that (iv) can be ingested by an existing freely-available prediction tool, `gnomonicus` ^8,12^. Together, this provides a transparent, data-driven framework for catalogue construction that is better suited to WGS-based surveillance.

## Methods

### Construction of the Training and Hold-out Validation Datasets

We downloaded v3.4.0 of the CRyPTIC dataset ^10^, containing 53,897 samples that have whole genome sequencing data with translated variant calls and at least one pDST result. This is referred to as the Entire Dataset, and the processing of the samples to produce this version of the CRyPTIC dataset has been described previously ^8^. Variation in the coding and promoter regions of each gene was detected and considered throughout the analysis; promoter regions were assumed to terminate after either 100 bases or when the next coding region was encountered, whichever came sooner. The CRyPTIC dataset also identifies genetic subpopulations (also referred to as minor alleles) supported by three or more short reads that have already passed quality control and have been classified as originating from MBTC ^8^. The pDST methods are varied; the two most common are 96-well broth microdilution plates (288,302 R/S measurements) and the BD Mycobacterial Growth Indicator Tube (190,909 R/S measurements). A set of published epidemiological cut-off (ECOFF) values ^13^ had been used to convert the MICs into R/S labels. Plate readings are subjective and we only took forward those that were concordant with a machine learning model, TMAS ^14^; this minimised exclusions to 159.

The Training Dataset (Table 1) was defined as all 41,130 samples found in v1.1.1 of the CRyPTIC dataset ^6^. This is identical to that used to build `WHOv1`^3^ but is missing seven samples; the reason for this is unknown. The remaining samples from v3.4.0 of the CRyPTIC dataset, which were collected and/or processed since the v1.1.1 release, formed The Validation Dataset (Table 1). It is likely that some of these samples were also used to build `WHOv2`.

**Table 1:** The number of samples, stratified by drug, in The Training Dataset (CRyPTICv1.1.1) and the hold-out Validation Dataset (CRyPTICv3.4.0 - CRyPTICv1.1.1). All samples have both drug susceptibility data and whole-genome sequences. Drugs are ordered as listed in `WHOv2`^2^. The first four compounds (RIF, INH, EMB and PZA) are the first-line treatment for drug-susceptible tuberculosis.

| **Drug** | **Samples in Training Dataset** |  |  | **Samples in Validation Dataset** |  |  |
|---|---|---|---|---|---|---|
|  | R | S | Total | R | S | Total |
| Rifampicin, RIF | 10,928 | 26,053 | 36,981 | 4,283 | 7,199 | 11,482 |
| Isoniazid, INH | 13,662 | 23,498 | 37,160 | 5,225 | 6,155 | 11,380 |
| Ethambutol , EMB | 5,525 | 28,980 | 34,505 | 2,033 | 5,003 | 7,036 |
| Pyrazinamide, PZA | 3,190 | 18,521 | 21,711 | 1,797 | 5,266 | 7,063 |
| Levofloxacin, LEV | 2,192 | 9,583 | 11,775 | 466 | 2,979 | 3,445 |
| Moxifloxacin, MXF | 2,983 | 14,207 | 17,190 | 829 | 6,576 | 7,405 |
| Bedaquiline, BDQ | 144 | 11,082 | 11,226 | 838 | 3,476 | 4,314 |
| Clofazimine, CFZ | 451 | 12,293 | 12,744 | 106 | 3,476 | 3,582 |
| Linezolid, LZD | 205 | 14,369 | 14,574 | 138 | 4,331 | 4,469 |
| Delamanid, DLM | 147 | 10,311 | 10,458 | 44 | 2,789 | 2,833 |
| Amikacin, AMI | 1,497 | 17,715 | 19,212 | 742 | 7,062 | 7,804 |
| Streptomycin, STM | 4,543 | 9,837 | 14,380 | 1,630 | 1,954 | 3,584 |
| Ethionamide, ETH | 2,887 | 11,596 | 14,483 | 1,053 | 3,806 | 4,859 |
| Kanamycin, KAN | 1,839 | 16,360 | 18,199 | 991 | 7,029 | 8,020 |
| Capreomycin, CAP | 956 | 10,582 | 11,538 | 595 | 5,461 | 6,056 |

### Identification of genetic variants statistically associated, or not associated, with resis- tance

We applied an algorithm originally termed ‘definitive defectives’ ^15^, later repurposed for generating MTBC catalogues ^16^, together with a two-tailed binomial test (Fig. S1), to identify and classify benign variants, as implemented in `catomatic` (v0.2.2) ^11,17^, as well as described and justified in the Supplement. This naturally leads to a ternary classification system: variants in resistance genes are classified as associated with resistance (for brevity, labelled ‘R’) or not associated with resistance (labelled ‘S’) or having an unknown effect (‘U’).

We focussed on genes implicated in conferring resistance because the univariate method can only use samples with a single mutation; including non-relevant genes reduces statistical power by limiting the number of informative isolates. Accordingly, and to enable a fair comparison to the performance of the WHO catalogues, we only considered the highly-confident (‘Tier 1’) candidate genes listed in `WHOv2` that contain at least one RAV ^5^. Synonymous and phylogenetic mutations (defined in *Merker et al.* ^18^) also reduce our power: all synonymous variants were excluded, and a small number of well-established phylogenetic variants (S95T, E21Q, G668D in *gyrA*, M291I in *gyrB*) were removed a priori as their prevalence across both resistant and susceptible isolates constrains the definitive defectives approach.

The background rate (*H*_0_), confidence level (*p*) and the minimum fraction of read support (FRS*_min_*) must also be selected - the latter is the minimum proportion of reads at a genetic loci required to support a variant call and is specified separately for building and applying catalogues. The process generated a catalogue (a CSV file) for each drug in a format that is compatible with `gnomonicus` ^8^ to maximise readability while enabling rapid application to new datasets.

### Evaluation and benchmarking of catalogue performance

When assessing catalogue performance, we predicted any sample containing a RAV for a particular drug to be Resistant (R) to that drug. If the sample contained only benign mutations, we predicted it to be Susceptible (S), whilst samples containing one or more variants with unknown effects (but no RAVs) were classified as Unknown (U). Since it is not reasonable to assume that all samples classified as having an Unknown effect are Susceptible for all drugs, we defined the Definite Prediction Rate (DPR) as the proportion of samples for which a definite prediction, (*n_R_* + *n_S_*)/(*n_R_*+ *n_S_* + *n_U_*), can be returned ^11^, where *n_R_* is the number of samples predicted to be resistant, etc. The sensitivity and specificity of the samples with a definite prediction (R or S) is then calculated. Usually one aims to minimise the very major error (VME) rate, which is the proportion of resistant samples incorrectly predicted to be susceptible (i.e. 1 *−* sensitivity), whilst also balancing the proportion of susceptible samples incorrectly predicted to be resistant, the major error (ME = 1 *−* specificity) rate. To enable us to algorithmically select a single parameter set (*H*_0_, *p*) per drug from the grid search (Fig. S2), we created a composite scoring function defined as a weighted sum of sensitivity (weight = 0*·*5), specificity (0*·*3), and DPR (0*·*2). Varying the weights either had no effect on the parameter set or, if it did, there was little effect on the subsequent performance of the resulting catalogue (Fig. S3).

To benchmark our catalogue against `WHOv1`, and to compare the performance of `WHOv1` to its reported results, we used a two-proportions Z-test on the Validation Dataset. These and any other reported p-values correspond to two-proportions Z-tests unless otherwise stated. To facilitate removal of the Expert Rules when comparing the algorithms the WHO ‘Final Confidence Grading’ was reverted to the ‘Initial Confidence Grading‘.

### Role of the funding source

The funders played no role in the design of this study.

## Results

### Setting parameters by drug improves performance

Anti-tuberculosis drugs act on a number of protein targets in MTBC and a range of resistance mechanisms have subsequently evolved. Accordingly, each drug’s catalogue should be built with tailored statistical parameters to maximise predictive performance. We therefore performed parameter grid searches on the Training Dataset using `catomatic` allowing the background threshold (0*·*05 *≤ H*_0_ *≤* 0*·*25) and confidence level (*p ∈* 0*·*9, 0*·*95) to vary independently for each drug (Fig. S2). Performance varied only slightly across the parameter space for most drugs, indicating a stable response surface. Drugs with many resistant samples in the Training Dataset, such as the first-line agents, consistently perform well on the Training Dataset, with the DPR, sensitivity, and specificity always exceeding 80%. In contrast, drugs with fewer resistant samples, including bedaquline, clofazimine, linezolid, and delamanid, exhibit both lower sensitivity (typically *<* 50%) and a stronger negative relationship with increasing background *H*_0_ value. For delamanid, the high proportion of unclassified variants also reduces the DPR. We programmatically selected parameters for each drug (Table S1) using a composite scoring function (Fig. S3) to rank candidate parameter sets, thereby creating our final catalogue. Rifampicin performs best, with a sensitivity of 96.1%, a specificity 98.5%, and a DPR 97.1%.

The number of variants associated or not associated with resistance varies by drug (Table 2) and gene (Table S2) and ranges from five variants classified for linezolid to 440 for pyrazinamide. Most genetic variants occur too infrequently to be classified, or are only observed in combination with others and thus remain unexposed to the algorithm, leaving many unclassified (Table S2); however, their combined impact is usually minimal as most resistance is typically (but not always) conferred by a few, highly ‘penetrant’ mutations. In the accompanying code repository and Supplement, this catalogue is versioned as CRyPTICv1.1.1-2025.8; throughout this manuscript, we refer to it as `catomatic-1`.

**Table 2:** The number of unique genetic variants by drug classified as resistance-associated (R) or not-associated (S) in the `catomatic-1` catalogue. Those unable to be classified i.e. have an unknown (U) phenotype are also listed. These have been stratified by gene in Table S2. The full drug names are given in Table 1

|  | RIF | INH | EMB | PZA | LEV | MXF | BDQ | CFZ | LZD | DLM | AMI | STM | ETH | KAN | CAP |
|---|---|---|---|---|---|---|---|---|---|---|---|---|---|---|---|
| R | 42 | 62 | 20 | 440 | 13 | 11 | 34 | 46 | 1 | 18 | 11 | 115 | 298 | 11 | 43 |
| S | 16 | 23 | 58 | 0 | 19 | 22 | 0 | 0 | 4 | 0 | 6 | 8 | 0 | 6 | 2 |
| Total | 58 | 85 | 78 | 440 | 32 | 33 | 34 | 46 | 5 | 18 | 17 | 123 | 298 | 17 | 45 |
| U | 352 | 626 | 1056 | 143 | 330 | 489 | 172 | 172 | 1 | 29 | 383 | 542 | 325 | 366 | 234 |

### Genetic subpopulations contribute to resistance and must be considered

Genetic sub-populations were detected in all key resistance genes. Visualising the prevalence of these minor alleles is complicated by (assumed) neutral lineage-associated variants that overwhelm the distribution, such as R463L in *katG*, as these are frequently observed in heterogenous samples. The gene *embB* harbours a disproportionately high number of benign minor alleles — 28.8% of all identified variants — in particular, R24P. Temporarily excluding these reveals that non-essential genes have a higher proportion of minor alleles compared with essential genes (8.91% vs 6.45%, *p <* 0.001, Fig. 1A), consistent with weaker negative selection constraints and a greater influence of genetic drift.

**Figure 1:**
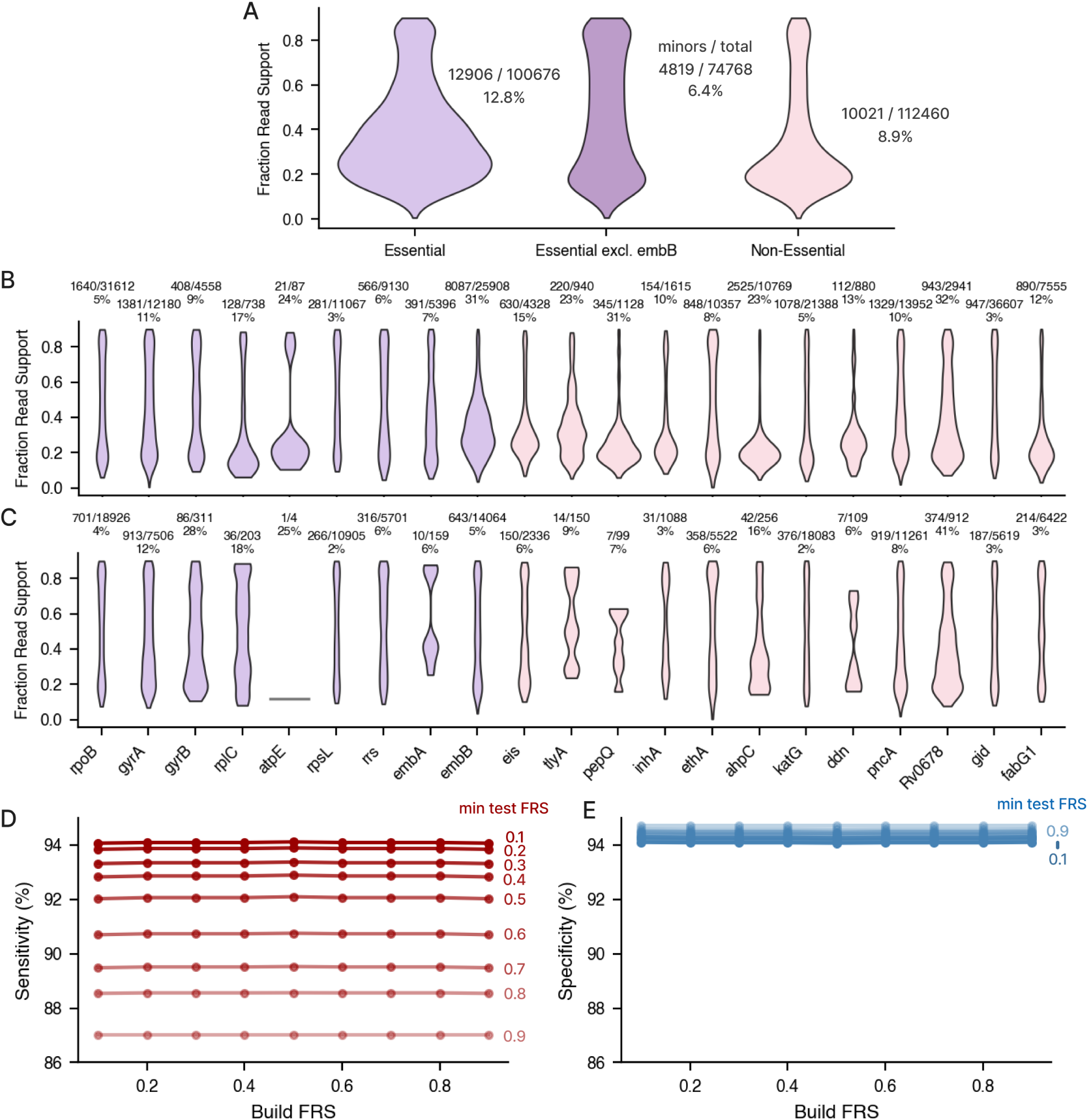
A. Distributions of the number of pooled minor alleles (*FRS <* 0.9) in The Training Dataset minus phylogenetic mutations, scaled against the total number of variants, from 0.1 *≤* FRS *<* 0.9 , for essential genes (lilac) and non-essential genes (pink). The dark purple violin excludes *embB*, whose minor alleles overwhelm the plot. The absolute fraction of minor alleles over the total number of alleles in The Training Dataset are shown - the area of each violin is proportional to this fraction. B. The number of minor alleles in The Training Dataset stratified by each candidate gene, scaled against the total number of variants observed for that gene, from 0.1 *≤* FRS *<* 0.9. C. The number of *known resistant* minor alleles in The Training Dataset stratified by each candidate gene, scaled against the total number of *known resistant* variants observed for that gene. We identify resistant alleles as those `catomatic` predicts to be RAVs when the minimum FRS is dropped to 0.10. The corresponding violins for *known non-resistance-associated* variants are in Fig. S4. D. Sensitivity and (E) specificity for moxifloxacin at varying FRS*_min_* thresholds used to train the catalogue (Build FRS) and when applying the catalogue (min test FRS). Complete sensitivity, specificity, and DPR results as a function of FRS*_min_* are shown for all drugs in Fig. S5.

Irrespective of essentiality, minor alleles are more likely to be found at a lower FRS of 10-40% (Fig. 1B), in contrast with the more even distributions observed for *resistant* sub-populations (Fig. 1C). Notably, the proportion of resistant samples that are heterogeneous exceeds 10% for several genes, including *gyrA* (12.2%), *gyrB* (27.7%), and *rplC* (17.7%), as well as the non-essential genes *Rv0678* (41.0%) and *ahpC* (16.4%).

Despite this, catalogues have historically been built assuming homogenous genetics, typically by requiring a high FRS*_min_*(0.75 or 0.9) to support variant calls, which also has the effect of mitigating short-read sequencing errors. To evaluate the impact of including these minor alleles, we built catalogues in the range 0.1 *≤* FRS*_min_ ≤* 0.9 and assessed their performance on the Training Dataset (Fig. S5). Lowering the FRS*_min_* when *building* the catalogues had a negligible impact on sensitivity or specificity (*<* 1%, *p >* 0.05, Table S3) for all drugs other than bedaquiline, clofazimine, and delamanid (whose catalogues perform poorly regardless). This suggests that resistance is usually conferred by mutations that can be identified and classified using a high FRS*_min_*, provided sufficient resistant samples are available.

A second FRS*_min_* threshold is set when *applying* the catalogue. Aside from linezolid and delamanid, lowering this FRS*_min_* increases the sensitivity for all drugs (*p <* 0.001, Table S3), enabling the detection of clinically relevant RAVs in subpopulations that would otherwise be missed. For example, the moxifloxacin catalogue built at FRS*_min_*= 0.1 is identical to that at FRS*_min_* = 0.9, yet sensitivity improves by seven percentage points when the detection threshold is lowered (Fig. 1D; Fig. S5; Table S3). Even for *rpoB*, which is highly essential and has a low incidence of minor RAVs (3.7%, Fig. 1C), excluding these variants from predictions decreases sensitivity by 2.1% (Table S3) ^19^. The largest corresponding increases in false positive predictions are observed for moxifloxacin (Fig. 1E) and ethambutol, for which specificity decreases by 0.63% (p = 0.027) and 0.50% (p = 0.032), respectively (Fig. S5, Table S3). The FRS*_min_* should therefore be lowered when *applying* the catalogue as this will significantly reduce the VME rate, at little cost to the ME rate on this dataset.

### Our automatically generated catomatic-1 catalogue performs similarly to WHOv1

Neither `WHOv1` nor `catomatic-1` were trained on the 14,380 samples in the Validation Dataset (Table 1) and therefore these can be used to assess their performance. The performance of `WHOv1`, when evaluated using a value for FRS*_min_* of 0.75, was largely consistent with that reported ^2^, indicating that `WHOv1` generalises well (Fig. S6, Table 3). We then evaluated the performance of `catomatic-1` on the Validation Dataset and compared its performance to `WHOv1` (Fig. 2, Table 3).

**Figure 2:**
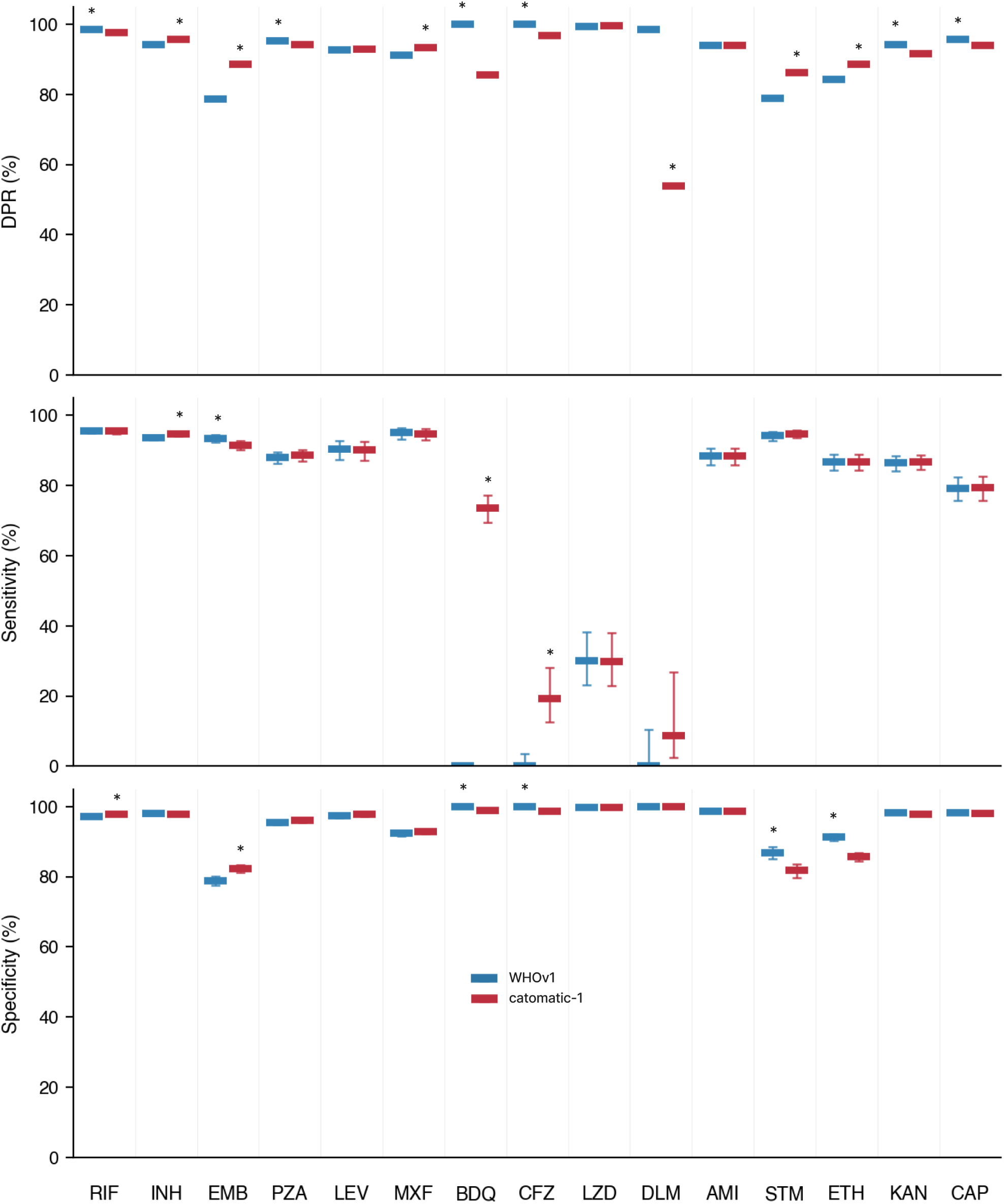
The definitive prediction rate (DPR), sensitivity and specificity, and for `WHOv1` (blue) and `catomatic-1` (red) catalogues when evaluated on The Validation Dataset of 14,380 samples using a minimum FRS of 0.1 to call genetic variants. Both `WHOv1` and `catomatic-1` were built on The Training Dataset and therefore these results are an independent assessment of performance. Asterisks show the significantly greater value at the 95% confidence level. Values are in Table 3.

**Table 3:** Sensitivity (Sens) (%), specificity (Spec) (%), and definite prediction rate (DPR) (%) for `catomatic-1` and `WHOv1` across drugs using the ternary prediction system, applied to The Validation Dataset. Values correspond to those plotted in Fig. 2 and are shown as point estimates with confidence intervals, computed from the reported lower and upper confidence limits as 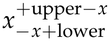. PPV and upper/lower bounds are in the Supplementary spreadsheet.

|  | catomatic-1 |  |  | WHOv1 |  |  |
|---|---|---|---|---|---|---|
| Drug | Sens | Spec | DPR | Sens | Spec | DPR |
| RIF | 95.5<sup>+0.59</sup><sub>−0.68</sub> | 97.9<sup>+0.31</sup><sub>−0.37</sub> | 97.6<sup>+0.27</sup><sub>−0.30</sub> | 95.5<sup>+0.58</sup><sub>−0.66</sub> | 97.1<sup>+0.36</sup><sub>−0.42</sub> | 98.6<sup>+0.20</sup><sub>−0.23</sub> |
| INH | 94.6<sup>+0.60</sup><sub>−0.66</sub> | 97.7<sup>+0.35</sup><sub>−0.42</sub> | 95.7<sup>+0.36</sup><sub>−0.39</sub> | 93.7<sup>+0.64</sup><sub>−0.71</sub> | 97.9<sup>+0.33</sup><sub>−0.40</sub> | 94.1<sup>+0.42</sup><sub>−0.45</sub> |
| EMB | 91.5<sup>+1.18</sup><sub>−1.35</sub> | 82.3<sup>+1.10</sup><sub>−1.16</sub> | 88.6<sup>+0.72</sup><sub>−0.76</sub> | 93.5<sup>+1.04</sup><sub>−1.22</sub> | 78.8<sup>+1.29</sup><sub>−1.35</sub> | 78.7<sup>+0.94</sup><sub>−0.97</sub> |
| PZA | 88.6<sup>+1.46</sup><sub>−1.64</sub> | 96.0<sup>+0.51</sup><sub>−0.58</sub> | 94.1<sup>+0.53</sup><sub>−0.57</sub> | 88.0<sup>+1.49</sup><sub>−1.68</sub> | 95.4<sup>+0.54</sup><sub>−0.61</sub> | 95.2<sup>+0.47</sup><sub>−0.53</sub> |
| LEV | 90.2<sup>+2.45</sup><sub>−3.14</sub> | 97.7<sup>+0.49</sup><sub>−0.63</sub> | 92.9<sup>+0.81</sup><sub>−0.91</sub> | 90.4<sup>+2.41</sup><sub>−3.11</sub> | 97.4<sup>+0.53</sup><sub>−0.67</sub> | 92.7<sup>+0.83</sup><sub>−0.91</sub> |
| MXF | 94.8<sup>+1.35</sup><sub>−1.79</sub> | 92.8<sup>+0.62</sup><sub>−0.67</sub> | 93.2<sup>+0.56</sup><sub>−0.59</sub> | 95.0<sup>+1.32</sup><sub>−1.76</sub> | 92.3<sup>+0.64</sup><sub>−0.71</sub> | 91.0<sup>+0.63</sup><sub>−0.67</sub> |
| BDQ | 73.6<sup>+3.71</sup><sub>−4.08</sub> | 98.8<sup>+0.32</sup><sub>−0.43</sub> | 85.5<sup>+1.02</sup><sub>−1.08</sub> | 0<sup>+0.46</sup><sub>−0</sub> | 100<sup>+0</sup><sub>−0.11</sub> | 100<sup>+0</sup><sub>−0.09</sub> |
| CFZ | 19.2<sup>+8.85</sup><sub>−6.54</sub> | 98.7<sup>+0.33</sup><sub>−0.44</sub> | 96.7<sup>+0.54</sup><sub>−0.64</sub> | 0<sup>+3.50</sup><sub>−0</sub> | 100<sup>+0</sup><sub>−0.11</sub> | 100<sup>+0</sup><sub>−0.11</sub> |
| LZD | 29.9<sup>+8.13</sup><sub>−7.04</sub> | 99.7<sup>+0.12</sup><sub>−0.22</sub> | 99.5<sup>+0.16</sup><sub>−0.24</sub> | 30.1<sup>+8.17</sup><sub>−7.08</sub> | 99.7<sup>+0.12</sup><sub>−0.22</sub> | 99.3<sup>+0.21</sup><sub>−0.29</sub> |
| DLM | 8.70<sup>+18.1</sup><sub>−6.28</sub> | 100<sup>+0</sup><sub>−0.25</sub> | 53.9<sup>+1.82</sup><sub>−1.84</sub> | 0<sup>+10.4</sup><sub>−0</sub> | 100<sup>+0</sup><sub>−0.14</sub> | 98.4<sup>+0.40</sup><sub>−0.53</sub> |
| AMI | 88.5<sup>+2.12</sup><sub>−2.53</sub> | 98.7<sup>+0.24</sup><sub>−0.30</sub> | 93.9<sup>+0.51</sup><sub>−0.56</sub> | 88.5<sup>+2.12</sup><sub>−2.53</sub> | 98.8<sup>+0.24</sup><sub>−0.30</sub> | 93.9<sup>+0.51</sup><sub>−0.55</sub> |
| STM | 94.8<sup>+1.03</sup><sub>−1.25</sub> | 81.8<sup>+1.81</sup><sub>−1.96</sub> | 86.1<sup>+1.10</sup><sub>−1.17</sub> | 94.2<sup>+1.14</sup><sub>−1.39</sub> | 86.8<sup>+1.61</sup><sub>−1.81</sub> | 78.9<sup>+1.30</sup><sub>−1.37</sub> |
| ETH | 86.8<sup>+2.07</sup><sub>−2.39</sub> | 85.7<sup>+1.13</sup><sub>−1.21</sub> | 88.5<sup>+0.86</sup><sub>−0.93</sub> | 86.7<sup>+2.09</sup><sub>−2.40</sub> | 91.2<sup>+0.93</sup><sub>−1.02</sub> | 84.2<sup>+1.00</sup><sub>−1.05</sub> |
| KAN | 86.8<sup>+1.98</sup><sub>−2.28</sub> | 97.7<sup>+0.34</sup><sub>−0.39</sub> | 91.7<sup>+0.58</sup><sub>−0.63</sub> | 86.4<sup>+2.01</sup><sub>−2.29</sub> | 98.2<sup>+0.29</sup><sub>−0.35</sub> | 94.2<sup>+0.49</sup><sub>−0.53</sub> |
| CAP | 79.3<sup>+3.17</sup><sub>−3.58</sub> | 98.1<sup>+0.34</sup><sub>−0.41</sub> | 93.9<sup>+0.58</sup><sub>−0.63</sub> | 79.3<sup>+3.18</sup><sub>−3.58</sub> | 98.1<sup>+0.33</sup><sub>−0.40</sub> | 95.6<sup>+0.49</sup><sub>−0.55</sub> |

Both catalogues achieved comparable performance across most drugs. Ethambutol showed the largest improvement over `WHOv1` (Table S5), with a 10% increase in DPR (88.6% v 78.6%, *p <* 0.001) and a +3.6% boost to specificity (82.3% v 78.8%, *p <* 0.001), balanced by a 2% reduction in sensitivity (91.5% v 93.5%, *p* = 0.021).

This reflects the classification of 47 variants not resistance-associated and eight RAVs in *embB* absent from `WHOv1`, partly offset by eight non-resistance-associated and three RAVs unique to `WHOv1` (Fig. 3). Isoniazid also performs better (*p <* 0.001), albeit the increases in DPR (95.7% v 94.1%) and sensitivity (94.6% v 93.7%) are more modest. Notably, `catomatic-1` makes fewer definite predictions for kanamycin and capreomycin (91.7% v 94.2% and 93.9 v 95.6%, respectively) than `WHOv1` and this is not offset by corresponding gains in sensitivity or specificity (Fig. 2, Table S5).

**Figure 3:**
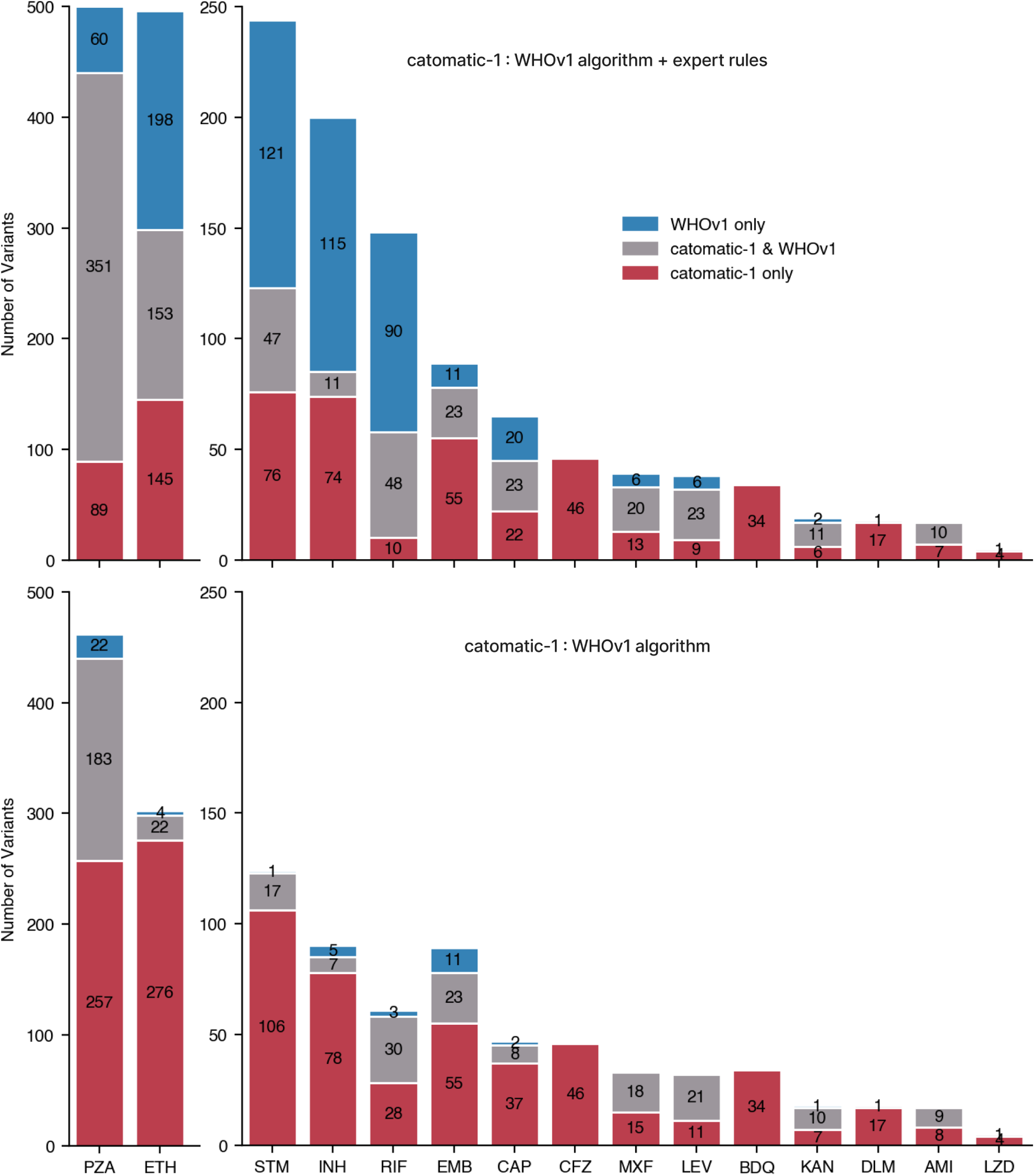
The number of unique genetic variants in The Training Dataset classified as either resistance-associated or non-associated differs between `catomatic-1` and `WHOv1`. Variants only classified by `catomatic-1` or `WHOv1` are shown in red and blue, respectively, whilst variants classified by both catalogues are shown in grey. (A) Naively comparing the two catalogues shows that, for many drugs, the number of variants that both catalogues classify is in the minority. (B) Removing the Expert Rules from `WHOv1` to enable a like-for-like comparison clearly shows how the more permissive statistics used by `catomatic` has allowed more definite classifications to be made. Bars for pyrazinamde (PZA) and ethionamide (ETH) are plotted at half scale for clarity. Unclassified mutations are not shown. Fig. S7 illustrates the effect each group has on predictive coverage. The full drug names are given in Table 1. Pairwise classification summaries are in Table S6, while per-variant classifications for `catomatic-1` and `WHOv1` with/without rules are summarised and compared in the supplementary spreadsheet

### Expert rules can improve performance, but limit end-to-end reproducibility

Although the performance of `WHOv1` and `catomatic-1` is similar, the number of genetic variants for which `WHOv1` returns a definite classification is greater due to generalisable rules which were added post-hoc to `WHOv1` by an Expert Panel for rifampicin, isoniazid, pyrazinamide, ethionamide, the fluoroquinolones, and the aminoglycosides ^2^. For example, an expert rule in `WHOv1` states that any genetic variant within the *rpoB* “Rifampicin Resistance Determining Region” are resistant and this allows `WHOv1` to classify 87 RAVs in the Training Dataset that `catomatic` does not, and 105 RAVs compared to its base algorithm; representing 71% of all mutations classified across both catalogues (Fig. 3A, S7A, Table S6). Conversely, `catomatic` also classifies a subset of rare variants that are not captured by the `WHOv1` algorithm or its expert rules.

Comparing `WHOv1`’s Initial and Final Confidence Grades shows that the net effect rules have on performance is small, particularly for rifampicin and isoniazid (Fig. S7B, Fig. S8), although they do increase the DPR values achieved by the `WHOv1` catalogue on the Validation Dataset for rifampicin, pyrazinamide, streptomycin, ethionamide (all *p <* 0.001), and amikacin (*p* = 0.030), albeit at costs to specificity (*p <* 0.01) for the first four drugs (Fig. S8, Table S4).

Excluding Expert Rules from `WHOv1` enables a like-for-like comparisons of the two statistical algorithms and demonstrates that the simpler statistical framework used by `catomatic` is more liberal, leading to many more *algorithmically* classified variants across all drugs compared to `WHOv1` (Fig. 3B, S7A, Table S6). The Expert Rules therefore improve the ability of `WHOv1` to make a prediction but at the price of requiring manual intervention and thus preventing the final catalogue from being end-to-end regenerated via a fully automated and reproducible process. We argue this constrains usability and reliability for a modest performance gain that could alternatively be achieved with adequate but less stringent statistical testing.

### Updating an algorithmically generated, version controlled catalogue is straightforward

We used `catomatic` to construct a final, larger catalogue on the Entire Dataset, versioned MBTC-CRyPTICv3.4.0-2025.8 and referred to here as `catomatic`-2. This catalogue cannot be independently validated and is presented to illustrate the effect of increased sample size on catalogue content. Incorporating the 14,380 additional samples (Table 1) enabled an additional 27 non-resistance-associated and 388 resistance-associated variants to be classified (Fig. S9, Supplementary spreadsheet), boosting the DPR of 9 drugs and increasing bedaquiline sensitivity by 10.3% (69.0% v 79.3%), bringing its performance close to a previously published catalogue ^11^ (Fig. S10, Table S7). We also calculated the performance of `WHOv1` and `WHOv2` on this dataset for completeness (Fig. S11, Table S7).

One must expect that when the evidence base is updated, classifications may change. For instance, the net gain in classified variants by `catomatic`-2 is 387 (not 415), as 28 mutations change classification. For example S441V in *rpoB* is no longer designated as an RAV since it now occurs in a susceptible isolate, while c-16t in the *embA* promoter, previously classified as susceptible, is now more observed in resistant isolates (34.6% vs 14.5%), resulting in a classification of associated with resistance (Fig. S9, Supplementary spreadsheet). Such changes are not only expected but are an intended feature of a statistical, surveillance-oriented approach, and can be readily tracked using version-controlled catalogues provided in a standardised, machine-readable format. However having variants change classification, as above, is undesirable when designing a molecular test, one of the use cases for catalogues. The obvious solution is to provide the statistical certainty and evidence alongside each RAV in the catalogue (as also done by `WHOv1`/v2) so classifications based on fewer samples (with therefore a greater probability of changing classification in future) can be screened out when using the catalogue in this way.

## Discussion

We have reproducibly built a comprehensive catalogue of genetic variants in *M. tuberculosis* associated with resistance to 15 anti-tuberculosis drugs (`catomatic-1`) using the same dataset that was used to build `WHOv1`^2^. Evaluating the performance of both catalogues on a hold-out Validation Dataset of 14,380 samples showed that, despite the process being automated and no Expert Rules being added, a simple statistical framework with reproducible, performance-optimised parameter selection can automatically detect and classify the majority of the predictive signal in the dataset, thereby achieving similar performance to `WHOv1` for all drugs.

Regardless of method, building a catalogue is more straightforward if (i) the resistance gene(s) is (are) essential (as this tends to lead to the resistance mechanisms being dominated by just a few genetic variants), (ii) variant effects are large (i.e. many-fold MIC increases), (iii) there is good concordance between different pDST methods, and there being (iv) many clinical samples to train on with (v) a high prevalence of resistance. These conditions are at least partially met here for most drugs, which is consistent with the generally satisfactory performance observed, except for clofazimine, delamanid and linezolid which all have fewer than 1,000 resistant samples in the Entire Dataset.

It is unsurprising that a significant proportion of MTBC complex samples show heterogeneous genetic variation, given that individuals may be infected for years before disease develops, allowing ample time for secondary infections and/or *in vivo* evolution ^19^. Heterogeneity is likely to be detected by WGS since it is usual to culture samples in liquid broth (MGIT tubes) and then extract DNA from multiple harvested ‘crumbs’. Rapidly and automatically building catalogues with different values of FRS*_min_* enabled us to examine the effect these subpopulations had on performance. Reducing FRS*_min_* had little effect when building catalogues, therefore we conclude most RAVs can be discovered and classified in homogenous samples (in this dataset at least). When we allowed minor alleles to be identified in samples, by lowering FRS*_min_*, the VME rate was reduced for all drugs (Fig. S5), consistent with other work ^1,5,11,20,21^. We observed improved performance down to an FRS*_min_* of 10%; ideally one would want to evaluate lower values but the CRyPTIC dataset contains very few minor alleles with an FRS*_min_ <* 10%, and secondly, the *optimal* threshold should be set statistically, but the CRyPTIC dataset lacks sufficient heterogeneous samples to permit this approach.

Like other approaches, our automated method based on `catomatic` requires users to specify the resistance-associated genes for each drug, meaning the resulting catalogue is only as reliable as this prior knowledge. This also does not encompass the regulatory networks in which these genes operate, and therefore is a simplified representation of the mechanisms of phenotypic resistance. A second assumption is that silent mutations and frequent phylogenetic variants (of which we filtered out four in the DNA gyrase) in the candidate genes have no effect. Using `catomatic`, one could perform a sensitivity analysis on the candidate genes and phylogenetic variants to justify their inclusion/exclusion.

The univariate, single-statistic approach preserves a direct mapping between the observed evidence and the classification of each variant, which is helpful in surveillance settings where results must facilitate auditable decision-making. However, it cannot capture combinatorial effects, including epistasis and compensation, nor can it implicitly determine lineage background effects or disentangle associations arising from clonal expansion, whereas phylogeny-aware multivariate regression in principle can ^20,22^. Moreover, dichotomising MIC distributions presupposes a clear bimodal separation between resistant and susceptible phenotypes, which is not always true, instead supporting the use of mixed-effects models trained directly on MICs ^22^.

Performance is key but should not come at the expense of usability, both when building and applying a catalogue. The recent proposal to use large language models to simplify interpreting the WHO catalogues ^23^ is testament to their unwieldiness; to build trust, catalogues *must* be easily readable by both human and computer. The catalogues output by `catomatic` are described using the GARC grammar ^8^ and can be stored in either JSON or CSV formats, both of which are readable using standard programming libraries, while CSVs can also be opened in spreadsheet software. The CSV files follow a simple tabular structure and are compatible with `gnomonicus` ^8,12^, a freely-available tool that returns a list of antibiotics and whether a sample is susceptible or resistant to each, and the effect of individual genetic variants, given a specified catalogue, a GenBank file and a variant call file.

The fast and reproducible method for generating catalogues demonstrated here, when coupled to a reliable genetic processing pipeline, could enable an international responsive surveillance system that in turn regularly and automatically updates a tuberculosis knowledge base. This would be particularly valuable for newly approved drugs (assuming they are accompanied by the rapid roll-out of pDST methods) by minimising the delay in detecting the first resistance-associated variants. As more samples are collected, catalogues could even be built using samples originating from a specific country or continent or by lineage, offering greater resolution in specific settings.

Enormous strides have brought us to this point, and it is vital we recognise that automation and reproducibility are both required to get us further and are also inevitable. By adopting best practises now we lay the groundwork for broader uptake and interoperability. This is feasible right now, and the ability to generate catalogues of tuberculosis resistance-associated genetic mutations at the press of a button is now a matter of will and implementation.

## Contributors

PWF, TMW, TEAP, DWC and ZI conceptualised the study with input from SO and NI. KMM, MH and DA identified additional samples to add to the Validation Dataset. JW, HT, MC, RDT and PWF processed all the samples creating the data tables. DA analysed these data, with supervision from PWF, DWE and TEAP. DA wrote the first draft of the manuscript which was reviewed by all authors prior to submission. All authors had full access to all the data in the study and had final responsibility for the decision to submit for publication.

## Declaration of interests

ZI, DWC & PWF receive consultancy fees from the Ellison Institute of Technology, Oxford Ltd.

## Data sharing

The versions of the CRyPTIC datasets used in this study are publicly available ^24^ and contain all phenotypic drug susceptibility testing measurements and all genetic variants detected in each sample; the datasets also includes the run accession numbers so the raw FASTQ files can be downloaded from the European Nucleotide Archive if required. An attendant, publicly accessible GitHub repository^*^ contains a script to download the relevant tables from v3.4.0 of the CRyPTIC dataset and Python code in the form of juypter notebooks that allows all results, including many of the figures, to be reproduced. The key dependency, `catomatic`, is installable via PyPI^†^.

## Acknowledgments

This research is funded by the National Institute for Health and Care Research (NIHR) Health Protection Research Unit in Healthcare Associated Infections and Antimicrobial Resistance (NIHR207397), a partnership between the UK Health Security Agency (UKHSA) and the University of Oxford, the National Institute for Health and Care Research Biomedical Research Centre: Oxford, the Ineos Oxford Institute for antimicrobial research and the Ellison Institute of Technology (EIT), Oxford Ltd. DA is supported by the EPSRC Sustainable Approaches to Biomedical Science: Responsible & Reproducible Research CDT and by ORACLE Corporation. The views expressed are those of the authors and not necessarily those of the NIHR, UKHSA, the Department of Health and Social Care or the Ellison Institute of Technology, Oxford Ltd. For the purpose of open access, the author has applied a CC BY public copyright licence to any Author Accepted Manuscript version arising from this submission.

## Footnotes

* https://github.com/fowler-lab/cryptic-catalogues-2025

† https://pypi.org/project/catomatic/

## Notes

https://github.com/fowler-lab/cryptic-catalogues-2025

### Competing Interest Statement

ZI, DWC & PWF work as consultants for the Ellison Institute of Technology, Oxford Ltd.

### Summary of Updates

Minor text changes and a new figure 1 based on feedback

