## Supplemental Information for "Rapidly and reproducibly building a comprehensive catalogue of resistance-associated variants for *M. tuberculosis* surveillance"

### Building resistance-associated genetic variant catalogues for *Mycobacterium tuberculosis* surveillance: an automated computational framework – Supplemental Information.

Dylan Adlard[1], Kerri M Malone[2], Jeremy Westhead[1], Martin Hunt[1,2], Hieu Thai[1], Matthew Colpus[1], Robert D Turner[1], Shaheed V Omar[3], David W Eyre[1,4,5], Nazir Ismail[6], Timothy M Walker[1], Timothy EA Peto[1,4,5], Derrick W Crook[1,4,5], Zamin Iqbal[2,7], and Philip W Fowler*[1,4,5]

[1]Nuffield Department of Medicine, John Radcliffe Hospital, University of Oxford, Headley Way, Oxford, U.K.

[2]European Bioinformatics Institute, Cambridge, UK

[3]Centre for Tuberculosis, National Institute for Communicable Diseases a Division of the National Health Laboratory Service, Johannesburg, South Africa

[4]Health Protection Research Unit in Healthcare Associated Infections and Antimicrobial Resistance, University of Oxford, Oxford, U.K.

[5]National Institute for Health and Care Research Biomedical Research Centre: Oxford, John Radcliffe Hospital, Headley Way, Oxford, UK

[6]Department of Clinical Microbiology and Infectious Diseases, University of the Witwatersrand, Johannesburg, South Africa

[7]Milner Centre for Evolution, University of Bath, UK

#### `catomatic's` classification algorithm

The exact method used to determine mutation effects is described in Adlard *et al.* (2024)[1]. A binomial test is used to classify benign mutations under the null hypothesis that resistance rates among carriers do not differ from expectation when the mutation is the *sole variant* in that drug's resistance genes. Resistance genes were defined as WHO tier-1 candidate genes with evidence of at least 1 resistance-associated variant (RAV)[2].

We assume resistance phenotypes occur by chance at a background proportion ($H_0$). A Wilson score interval with a confidence level $p$ is used to determine if there is a predominance of susceptible samples carrying that mutation below this value (Fig. S1). If significant, this mutation is labelled not resistance-associated (S, i.e. benign) and masked from training to potentially proffer additional informative mutations. This relies on the assumption that mutations which do not confer resistance individually also do not contribute when co-occurring with RAVs.

Once all benign mutations have been identified and masked, the binomial test is reapplied at confidence $p$ to determine if samples containing each remaining mutation are observed at a proportion of resistance greater than ($H_0$), resulting in a resistance-associated (R) classification. Mutations not reaching significance in either direction, often due to limited sample size, are reported as uncertain (U), yielding a ternary classification (R/S/U).

This formulation differs from approaches based on odds ratios and composite criteria in that it directly models the conditional probability of resistance given a mutation, $p(R \mid M)$. In contrast, odds ratios quantify the relative association between mutation and phenotype by comparing carriers to a background population of non-carriers, $p(R \mid M)$ relative to $p(R \mid \neg M)$, and are therefore sensitive to the composition of the non-carrier group. Furthermore, the binomial formulation provides well-behaved estimates and confidence intervals even in sparse data regimes with small or zero cell counts.

By conditioning only on isolates carrying the mutation, the binomial test avoids reliance on a heterogeneous control group and yields a direct mapping from observed phenotype counts to classification. Importantly, this framework naturally supports a ternary classification (R/S/U), allowing uncertainty arising from limited sample size to be explicitly represented in a single step. This facilitates transparency, in which each assignment can be immediately traced to the underlying data. We prioritise this direct estimation of per-mutation resistance probabilities, while acknowledging that this approach does not model interactions between mutations or other higher-order effects.

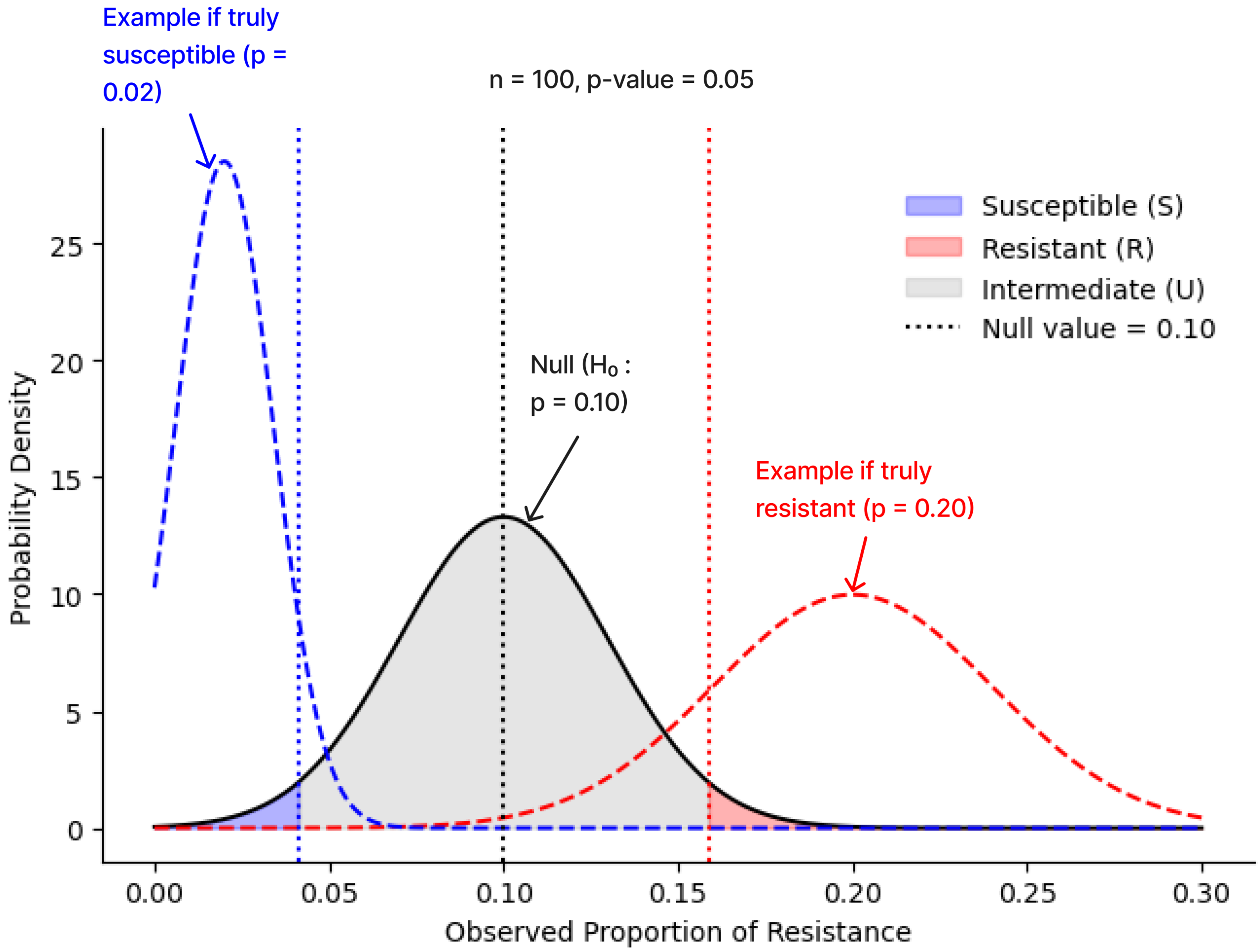


Figure S1: Illustration of resistance classification using a two-tailed binomial test (normal approximation, n = 100, chosen threshold p = 10%, confidence interval = 95%). The black curve represents the sampling distribution of observed resistance proportions under the null hypothesis that the mutation has no effect (ie, resistance rate = 10%). Observed proportions are classified as not-associated (S) if significantly lower than the null (blue shaded region), resistance-associated (R) if significantly higher (red shaded region), and Unclassified (U) if not significantly different (grey region). Dashed red and blue curves are hypothetical distributions for mutations truly associated with resistance (p=0.20) and susceptibility (p = 0.02), respectively. The degree of overlap between true-effect distributions and the null represents the power of the test.

| Drug | Background Rate | Confidence interval | Training Performance (%) Sensitivity | Specificity | DPR |
|---|---|---|---|---|---|
| RIF | 0.20 | 0.90 | 96.1 | 98.5 | 97.1 |
| INH | 0.25 | 0.90 | 94.6 | 98.8 | 95.7 |
| EMB | 0.25 | 0.90 | 94.7 | 92.0 | 90.8 |
| PZA | 0.05 | 0.90 | 86.4 | 97.3 | 97.6 |
| LEV | 0.25 | 0.90 | 92.3 | 97.1 | 93.3 |
| MXF | 0.25 | 0.95 | 94.1 | 94.1 | 93.2 |
| BDQ | 0.05 | 0.90 | 52.2 | 99.0 | 95.8 |
| CFZ | 0.05 | 0.90 | 19.5 | 98.7 | 96.4 |
| LZD | 0.25 | 0.95 | 29.8 | 99.8 | 99.7 |
| DLM | 0.05 | 0.90 | 34.1 | 99.9 | 61.6 |
| AMI | 0.05 | 0.90 | 87.4 | 98.3 | 92.1 |
| STM | 0.15 | 0.90 | 95.4 | 93.2 | 91.5 |
| ETH | 0.05 | 0.90 | 90.9 | 87.0 | 89.6 |
| KAN | 0.10 | 0.90 | 83.6 | 97.1 | 91.8 |
| CAP | 0.05 | 0.90 | 84.0 | 97.6 | 92.3 |

Table S1: Optimal parameter settings as chosen by the grid search for each drug, and the sensitivity, specificity, and Definite Prediction Rate (DPR) per drug in the `catomatic-1` catalogue when applied to the training set. All performance values are percentages, rounded to three significant figures. Optimal parameter settings were selected using a weighted composite score prioritising sensitivity, followed by specificity and DPR (see Methods).

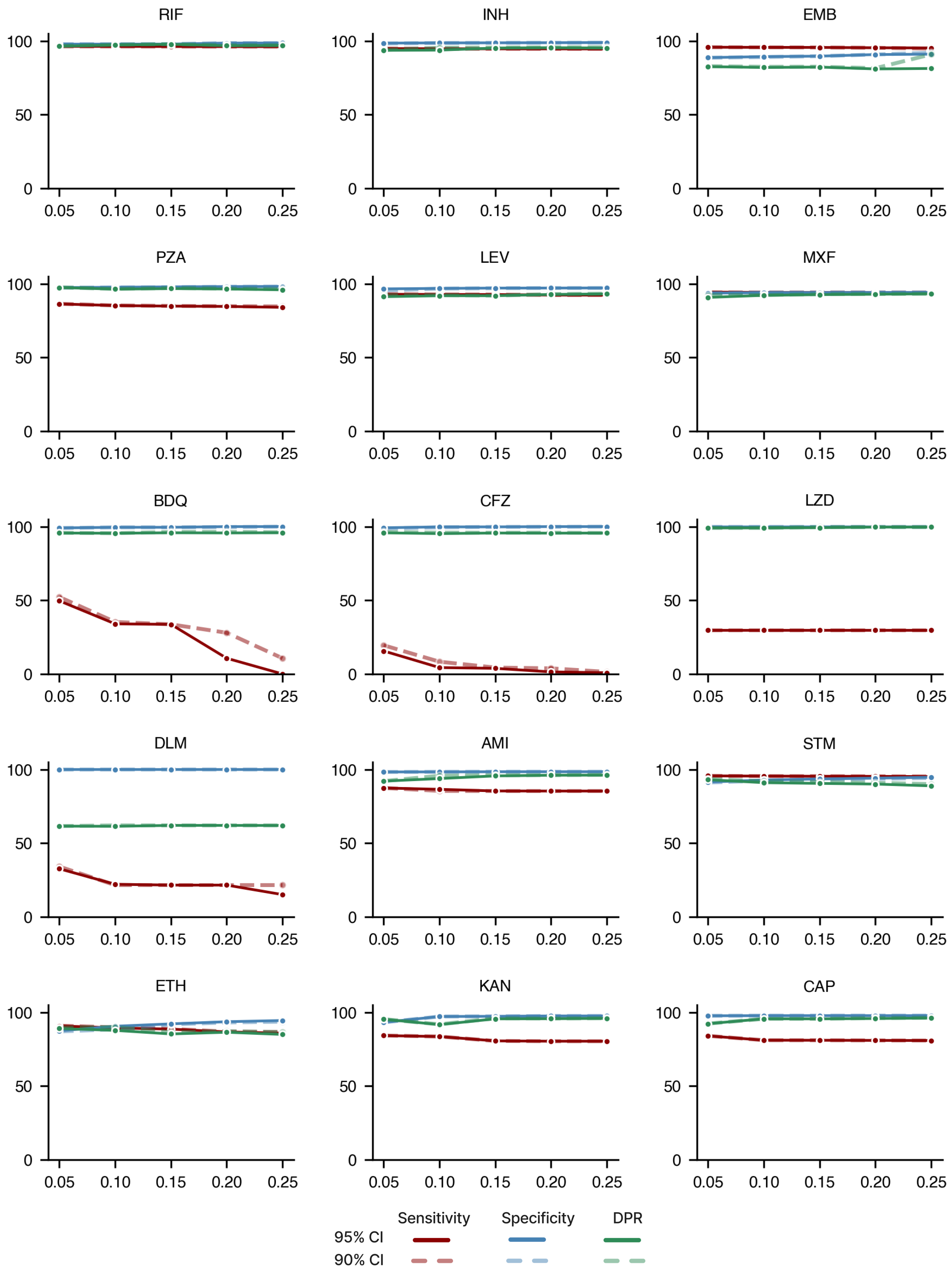


Figure S2: Sensitivity, specificity, and DPR for catalogues trained on varying background rates (0.05 increments) and at two confidence levels, 95% (solid lines) and 90% (dotted lines), when applied to the Training Dataset only. For drugs where there is limited resistant data (BDQ, DLM, LZD, and CFZ), raising the background rate and to a minor extent, the p-value, is detrimental to sensitivity and hence lower values of 0.05 and 0.90 are preferable, respectively.

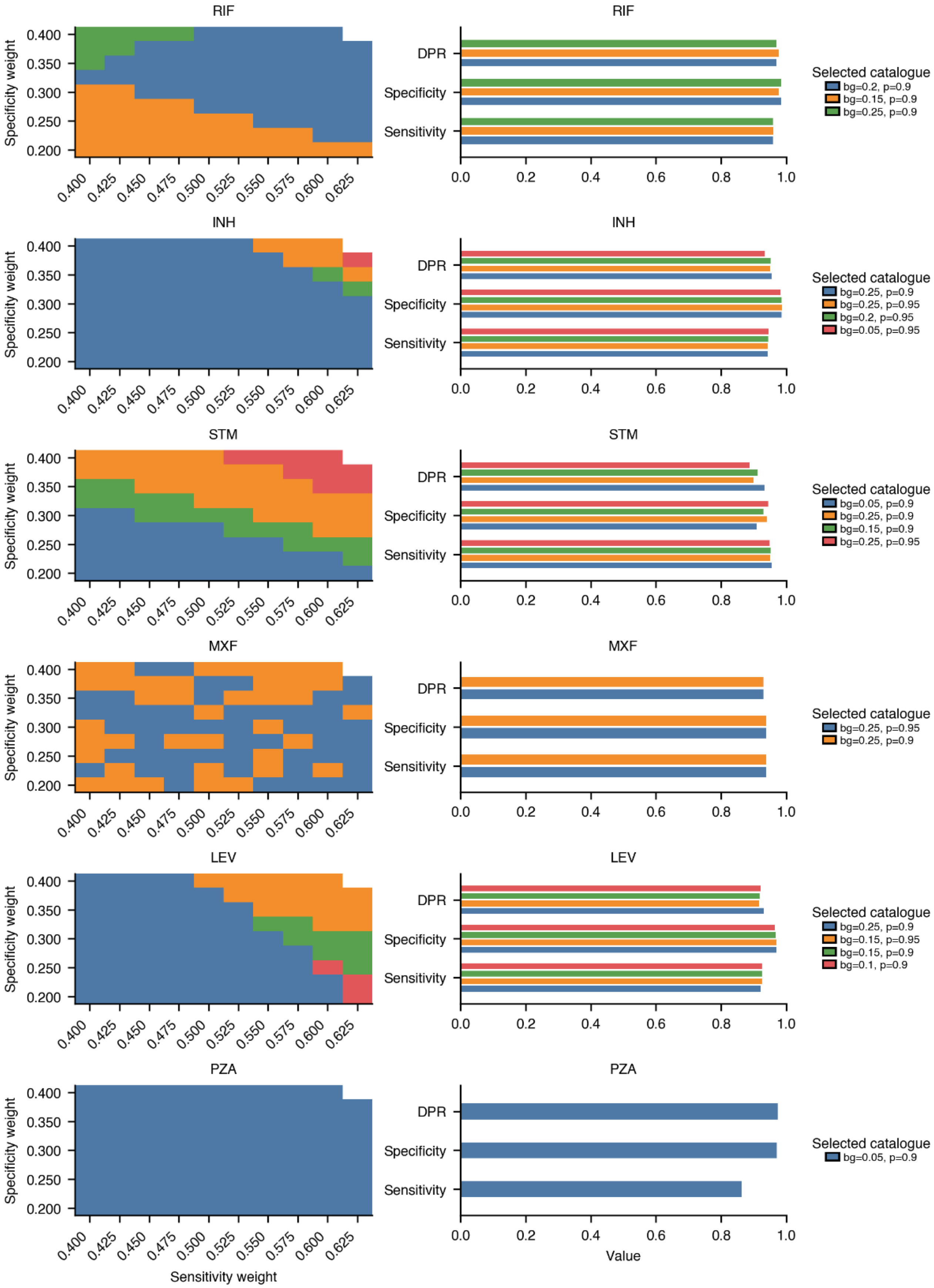
RIF
INH
STM
MXF
LEV
PZA
Specificity weight
0.400
0.350
0.300
0.250
0.200
0.400 0.425 0.450 0.475 0.500 0.525 0.550 0.575 0.600 0.625
Sensitivity weight
DPR
Specificity
Sensitivity
0.0 0.2 0.4 0.6 0.8 1.0
Value
Selected catalogue
bg=0.2, p=0.9
bg=0.15, p=0.9
bg=0.25, p=0.9
Selected catalogue
bg=0.25, p=0.9
bg=0.25, p=0.95
bg=0.2, p=0.95
bg=0.05, p=0.95
Selected catalogue
bg=0.05, p=0.9
bg=0.25, p=0.9
bg=0.15, p=0.9
bg=0.25, p=0.95
Selected catalogue
bg=0.25, p=0.95
bg=0.25, p=0.9
Selected catalogue
bg=0.25, p=0.9
bg=0.15, p=0.95
bg=0.15, p=0.9
bg=0.1, p=0.9
Selected catalogue
bg=0.05, p=0.9

Figure S3

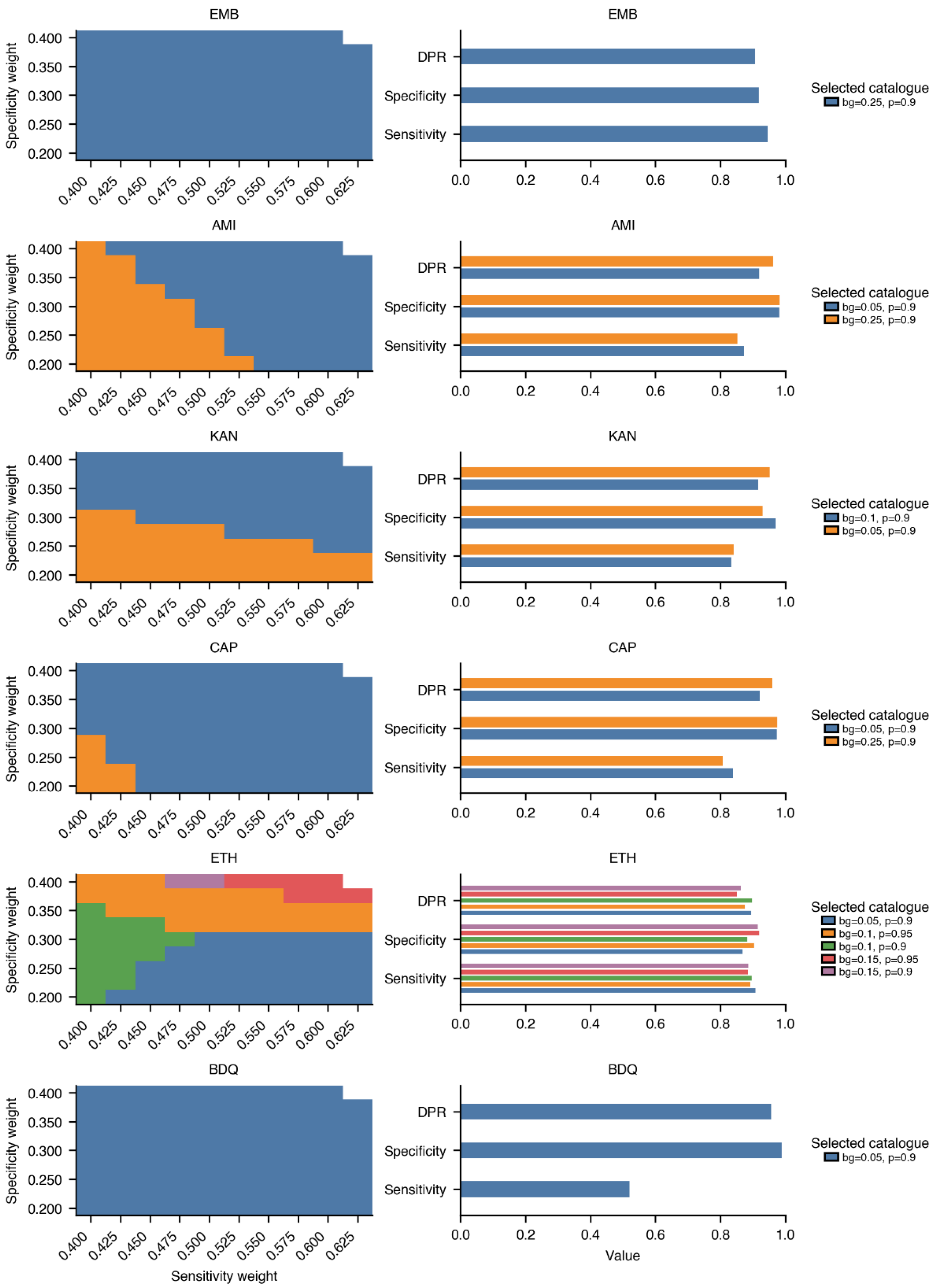
EMB
AMI
KAN
CAP
ETH
BDQ
Specificity weight
Sensitivity weight
DPR
Specificity
Sensitivity
Value
Selected catalogue
bg=0.25, p=0.9
bg=0.05, p=0.9
bg=0.25, p=0.9
bg=0.1, p=0.9
bg=0.05, p=0.9
bg=0.05, p=0.9
bg=0.25, p=0.9
bg=0.05, p=0.9
bg=0.1, p=0.95
bg=0.1, p=0.9
bg=0.15, p=0.95
bg=0.15, p=0.9
bg=0.05, p=0.9

Figure S3

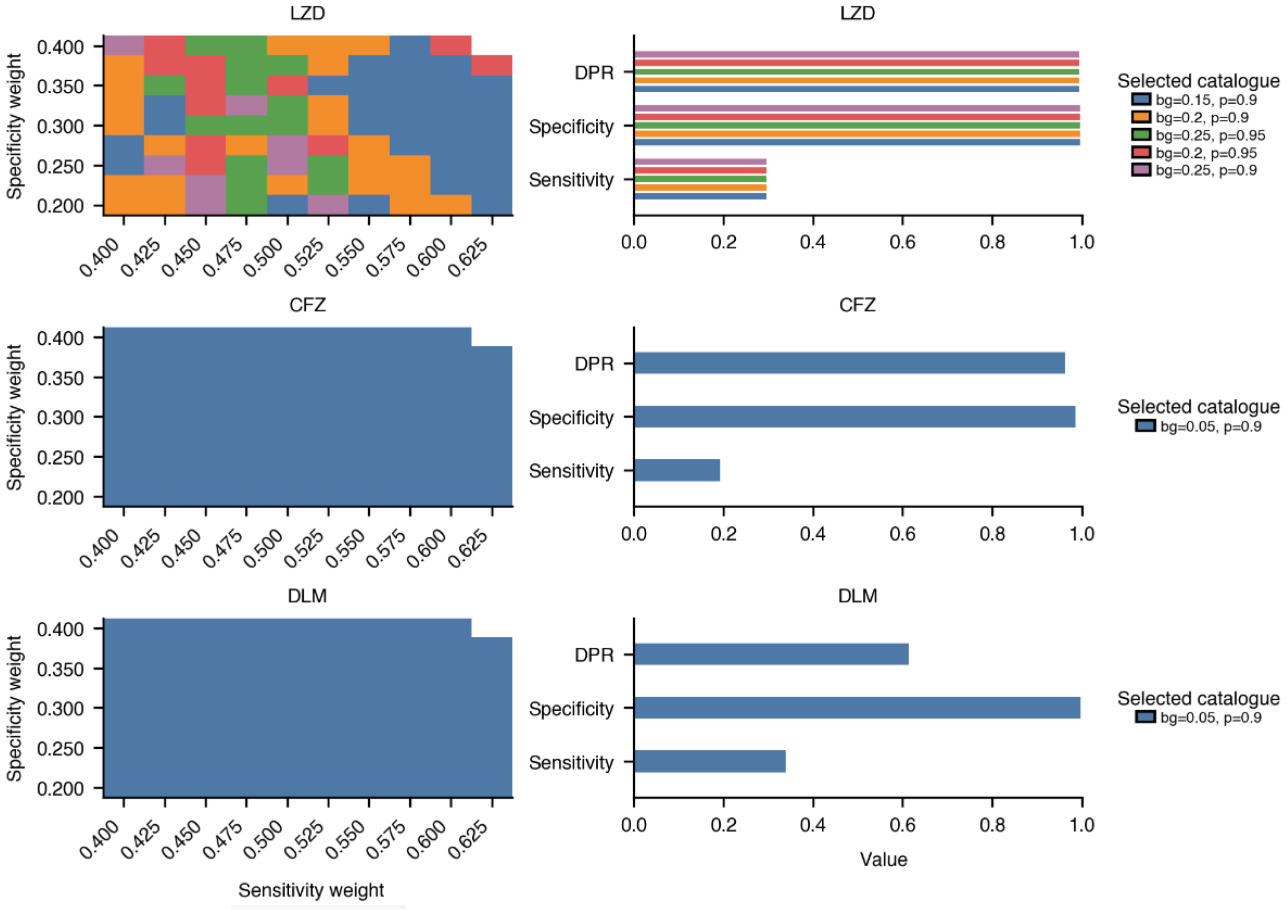


Figure S3: Catalogue selection response to varying weights in the score function. For each drug, the left panel shows the catalogue selected as optimal across a grid of weightings applied to sensitivity, specificity, and definite prediction rate (DPR), with colours indicating different parameter sets. The right panel shows the corresponding performance of each selected catalogue. For most drugs, a single or pair of catalogues is consistently selected across weightings, indicating robust parameter choice; where variation occurs, the resulting differences in performance metrics are minimal and largely confined to drugs with weaker signal.

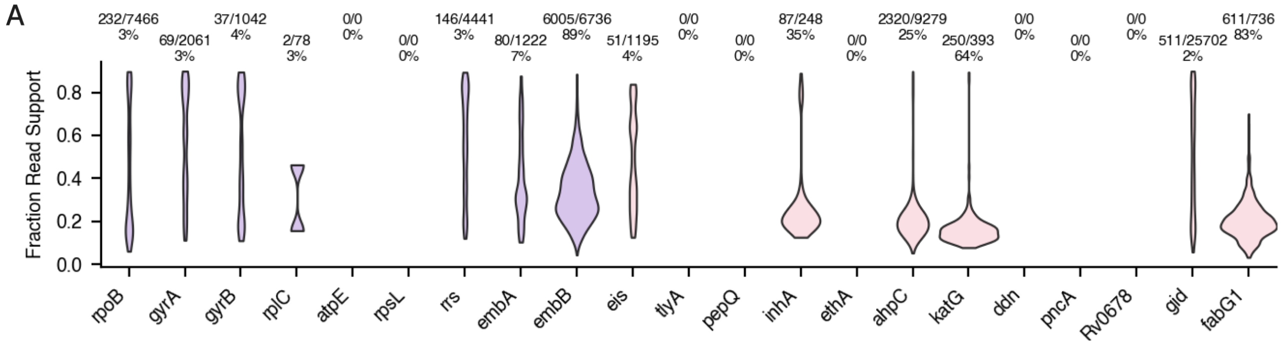


Figure S4: A. The number of *known non-resistance-associated* minor alleles in The Training Dataset stratified by each candidate gene, scaled against the total number of *known non-associated* variants observed for that gene. This corresponds to Fig. 1 in the main text. We identify non-associated alleles as those `catomatic` predicts to be benign when the minimum FRS is dropped to 0.10. There are gaps because we have not catalogued benign variants for those genes. Equally, these numbers and those in Fig. 1C do not add up to Fig. 1B, because Fig. 1B contains *all* variants, not just those for which we have determined the effect. The Essential genes are coloured lilac and non-essential are pink.

| Drug | Gene | R | S | Total | U observed | U unobserved |
|---|---|---|---|---|---|---|
| RIF | *rpoB* | 42 | 16 | 58 | 352 | 648 |
| INH | *ahpC* | 3 | 7 | 10 | 96 | 110 |
| | *fabG1* | 5 | 4 | 9 | 52 | 60 |
| | *inhA* | 4 | 5 | 9 | 52 | 64 |
| | *katG* | 50 | 7 | 57 | 426 | 776 |
| EMB | *embA* | 2 | 30 | 32 | 520 | 376 |
| | *embB* | 18 | 28 | 46 | 536 | 363 |
| PZA | *pncA* | 440 | 0 | 440 | 143 | 536 |
| LEV | *gyrA* | 10 | 13 | 23 | 187 | 630 |
| | *gyrB* | 3 | 6 | 9 | 143 | 475 |
| MXF | *gyrA* | 9 | 15 | 24 | 282 | 534 |
| | *gyrB* | 2 | 7 | 9 | 207 | 411 |
| BDQ | *Rv0678* | 30 | 0 | 30 | 103 | 402 |
| | *atpE* | 1 | 0 | 1 | 8 | 32 |
| | *pepQ* | 3 | 0 | 3 | 61 | 172 |
| CFZ | *Rv0678* | 40 | 0 | 40 | 104 | 391 |
| | *pepQ* | 6 | 0 | 6 | 68 | 162 |
| LZD | *rplC* | 1 | 4 | 5 | 29 | 94 |
| DLM | *ddn* | 15 | 0 | 15 | 45 | 146 |
| | *dprE1* | 3 | 0 | 3 | 62 | 150 |
| AMI | *eis* | 4 | 3 | 7 | 138 | 278 |
| | *rrs* | 7 | 3 | 10 | 245 | 394 |
| STM | *gid* | 102 | 5 | 107 | 369 | 608 |
| | *rpsL* | 5 | 0 | 5 | 20 | 65 |
| | *rrs* | 8 | 3 | 11 | 153 | 485 |
| ETH | *ethA* | 283 | 0 | 283 | 276 | 957 |
| | *fabG1* | 7 | 0 | 7 | 28 | 86 |
| | *inhA* | 8 | 0 | 8 | 21 | 96 |
| KAN | *eis* | 7 | 2 | 9 | 130 | 284 |
| | *rrs* | 4 | 4 | 8 | 236 | 405 |
| CAP | *rrs* | 5 | 2 | 7 | 170 | 472 |
| | *tlyA* | 38 | 0 | 38 | 64 | 243 |

Table S2: Per-gene mutation counts for each drug in `catomatic-1`. Totals reflect only R and S counts. `U observed` corresponds to mutations exposed to algorithm but with insufficient power to be classified, while `U unobserved` variants never occur alone and therefore were never exposed.

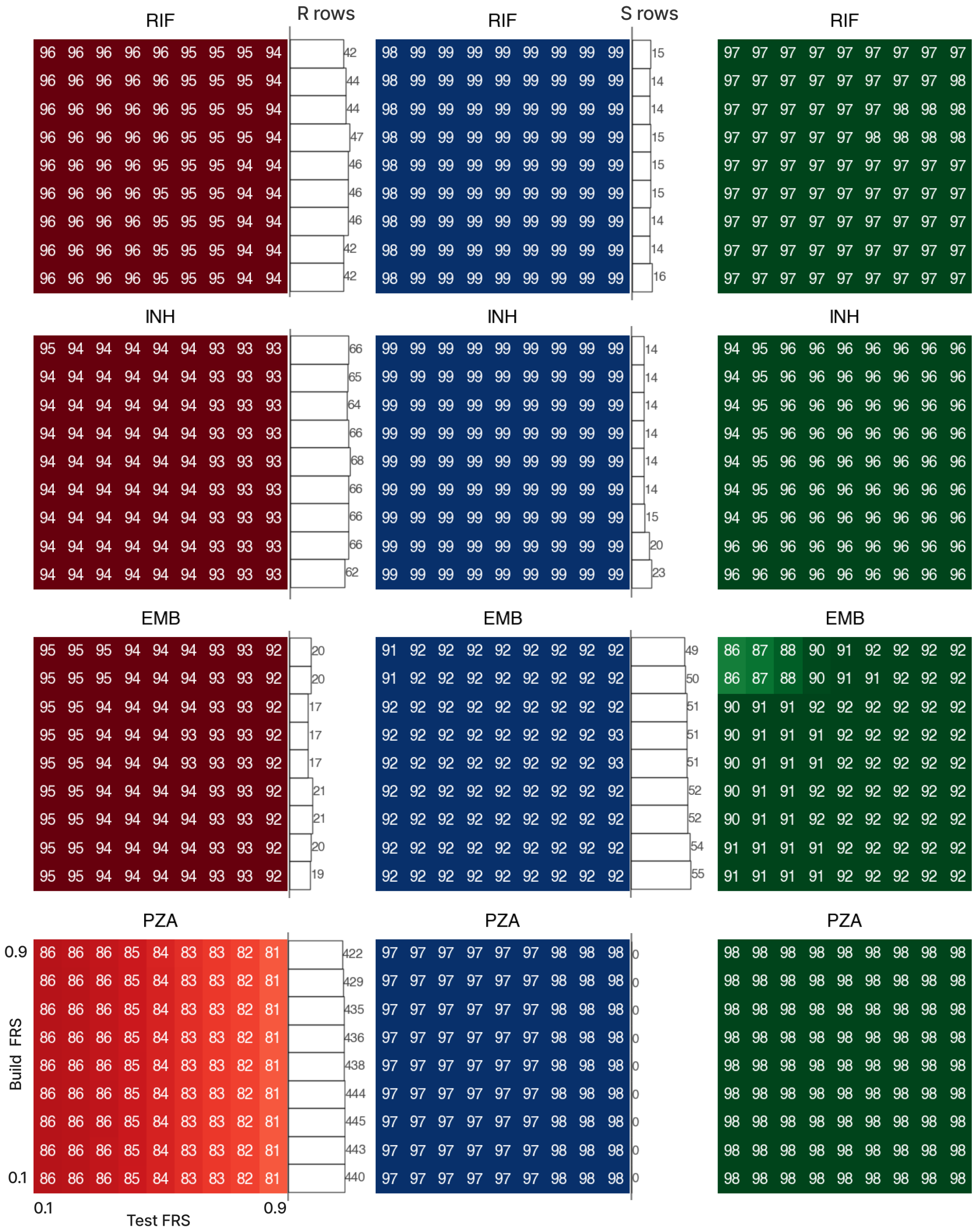


Figure S5

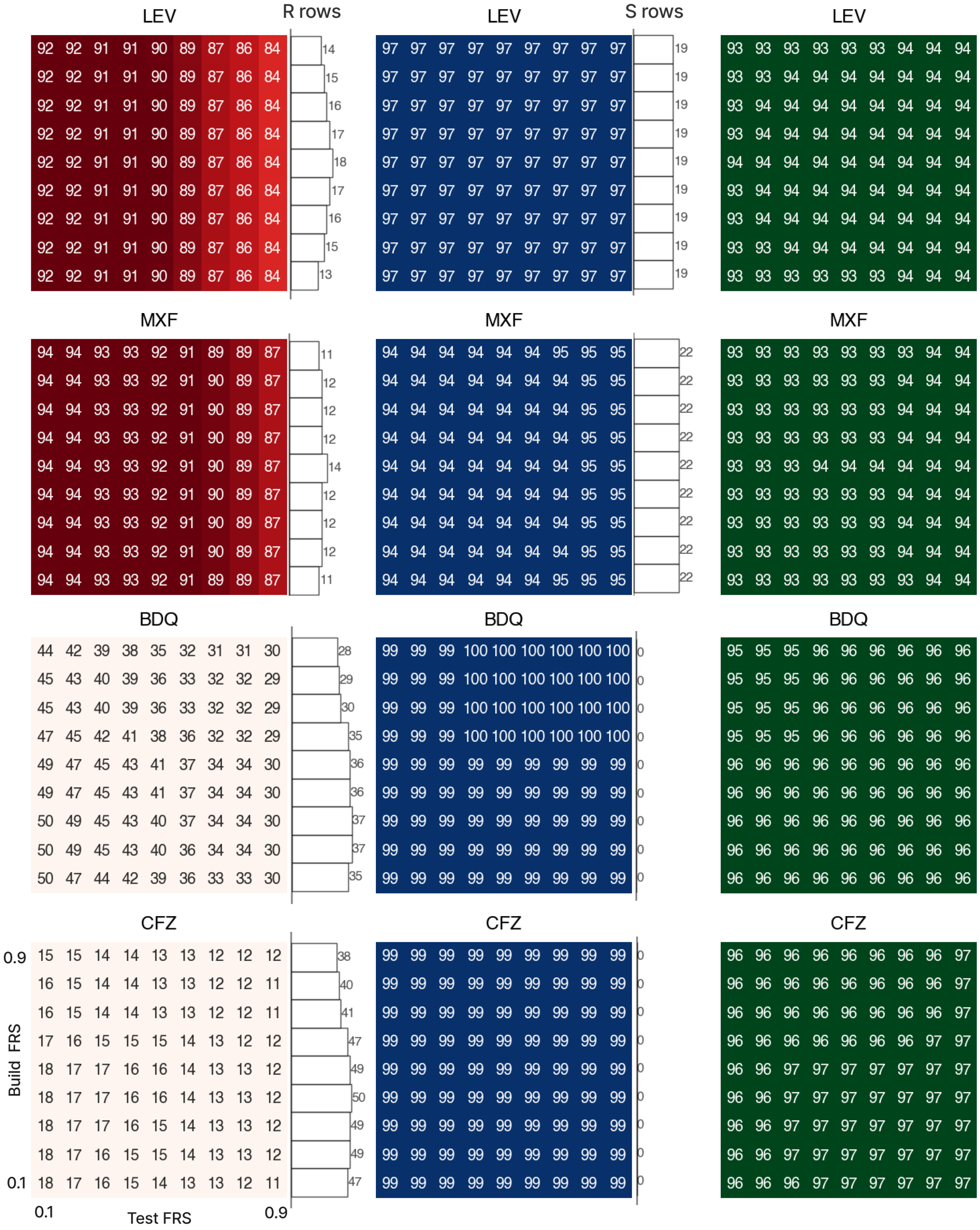
LEV
R rows
LEV
S rows
LEV
MXF
MXF
MXF
BDQ
BDQ
BDQ
CFZ
CFZ
CFZ
0.9
0.1
Build FRS
0.1
Test FRS
0.9

Figure S5

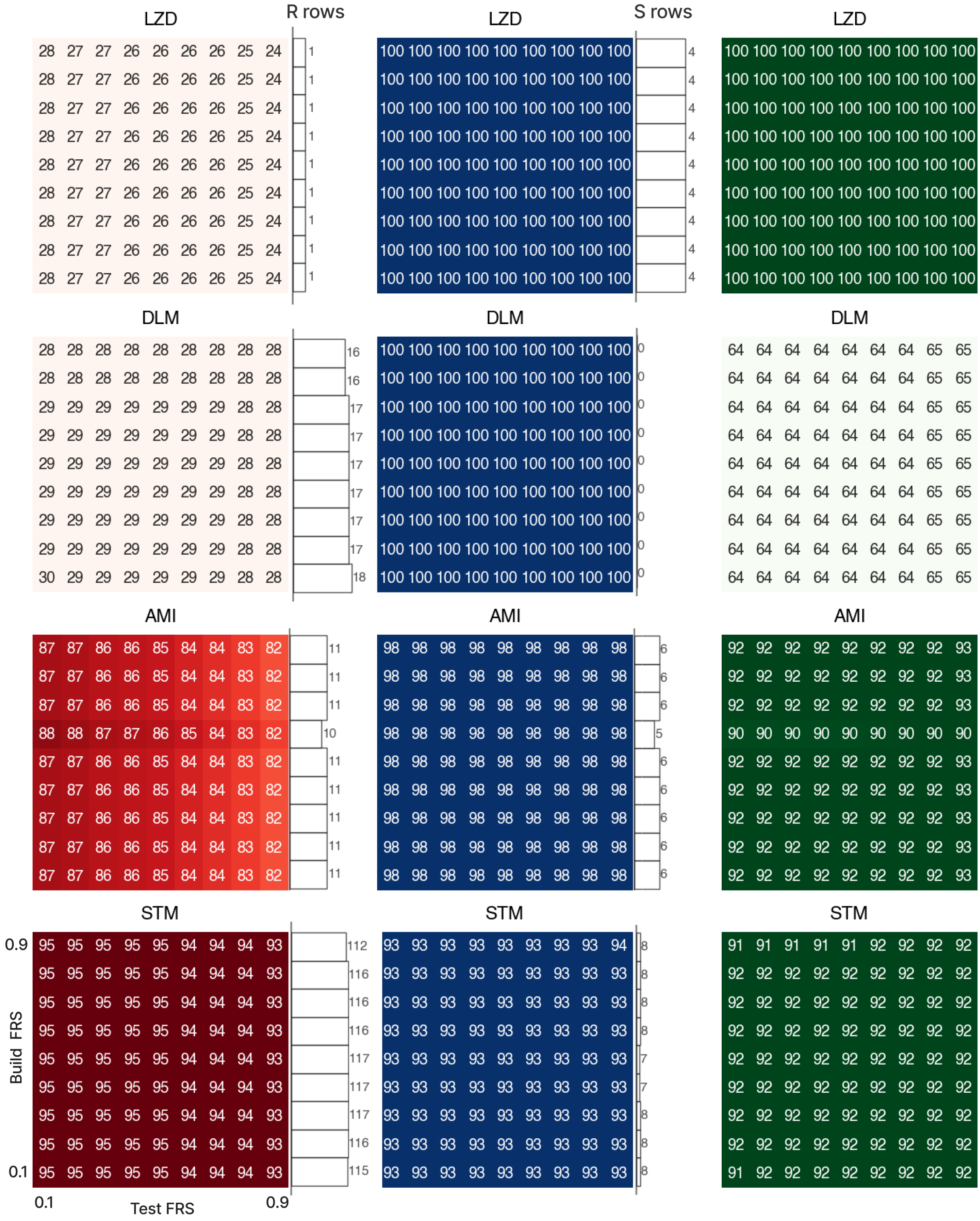


Figure S5

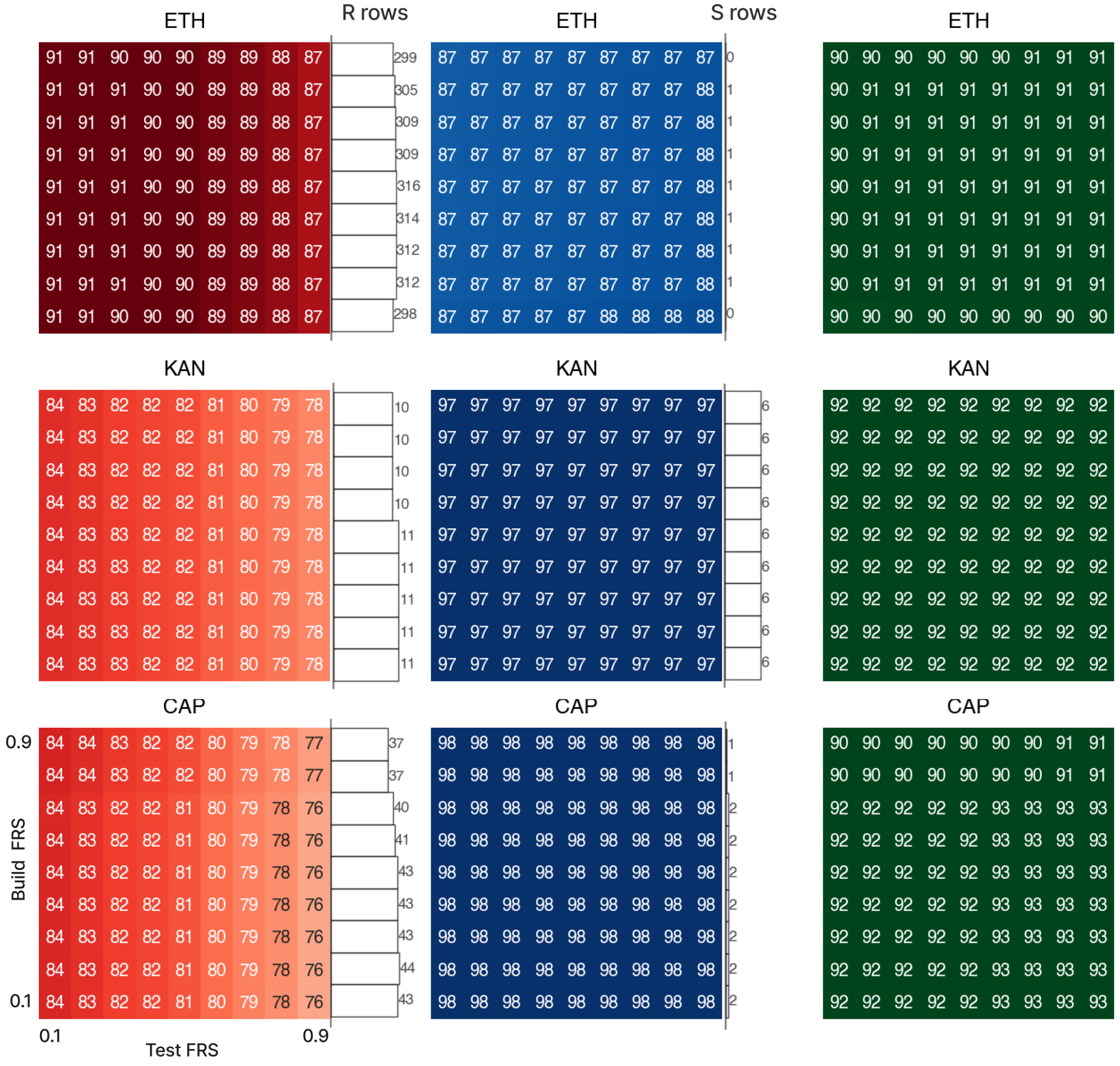


Figure S5: Heatmaps of sensitivity (red), specificity (blue), and DPR (green) for catalogues built and validated at different FRS values on The Training Dataset, for all drugs. The number of resistance associated (R) catalogued for each Build FRS is plotted on the right of each sensitivity heatmap, and the number of not resistance-associated (S) variants plotted on the right of each specificity heatmap. Differences in performance metrics and associated p-values when building and testing at $FRS_{min}$ are shown in Table S3. It is unsurprising that lowering the $FRS_{min}$ affects how many variants are classified, as many rare variants are only observed at lower FRS values in this dataset. Because the univariate classification algorithm is sensitive to the set of informative, exposed mutations, changes in this threshold can also lead to local fluctuations in performance, such as the increase observed for amikacin at $FRS_{min}$ = 0.6, where a previously misclassified variant is not accessible to the algorithm. Similarly, lowering the $FRS_{min}$ when building the catalogue *does not* always *increase* the number of classified variants, as seen for several drugs; however, because these variants are rare, the impact on sensitivity is typically very low (Table S3). Per-variant classifications at the $FRS_{min}$ build values are compared in the supplementary spreadsheet.

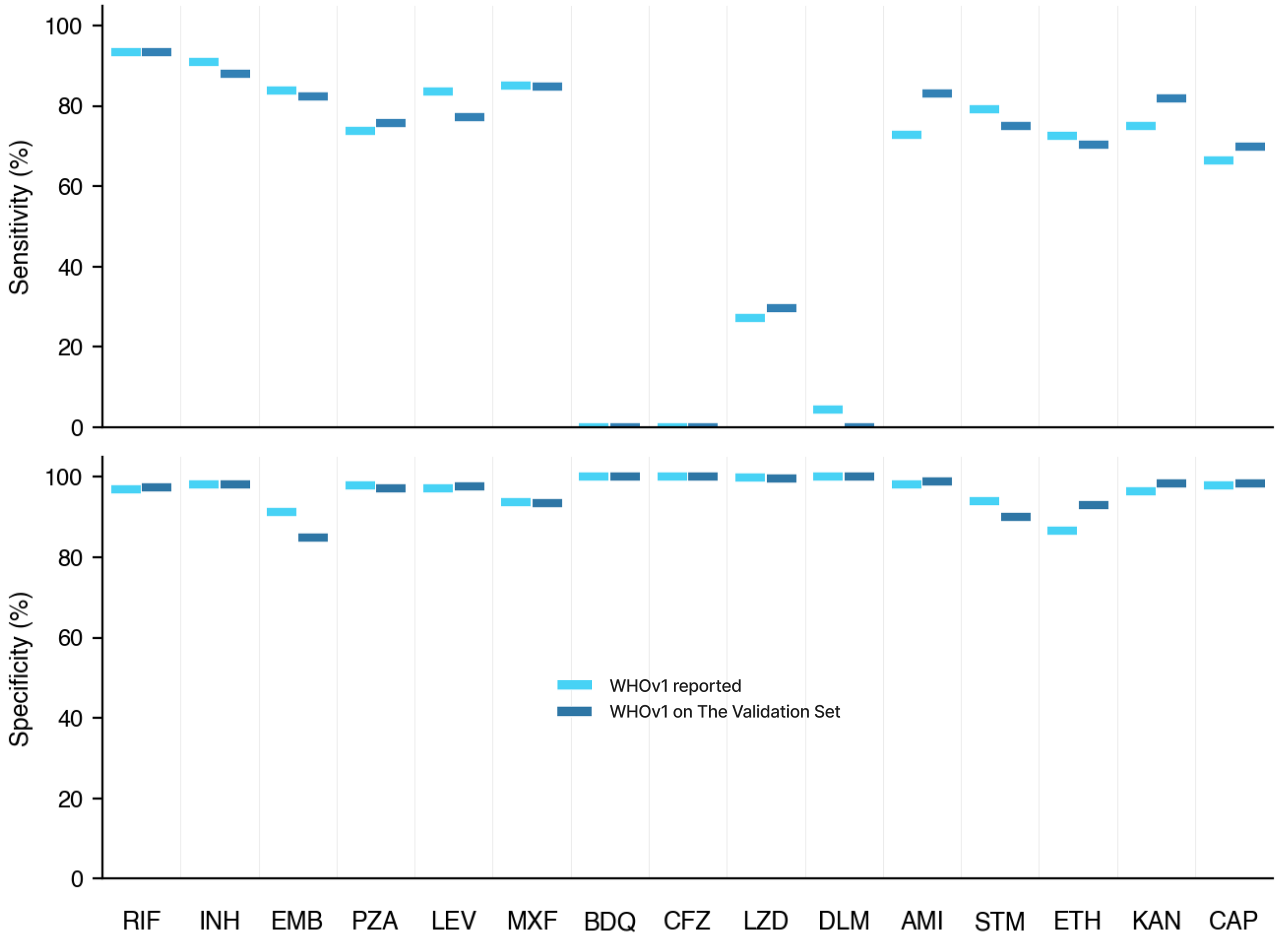


Figure S6: The sensitivity and specificity of the `WHOv1` catalogue (light blue) reported in the official WHO documentation[3], and the performance of WHOv1 when evaluated on the The Validation Dataset of 14,380 samples (dark blue) using an $FRS_{min}$ of 0.75. `WHOv1` has not seen samples in The Validation Dataset, providing a more fair estimate of performance. No variants associated with resistance to CFZ or BDQ were reported in `WHOv1`.

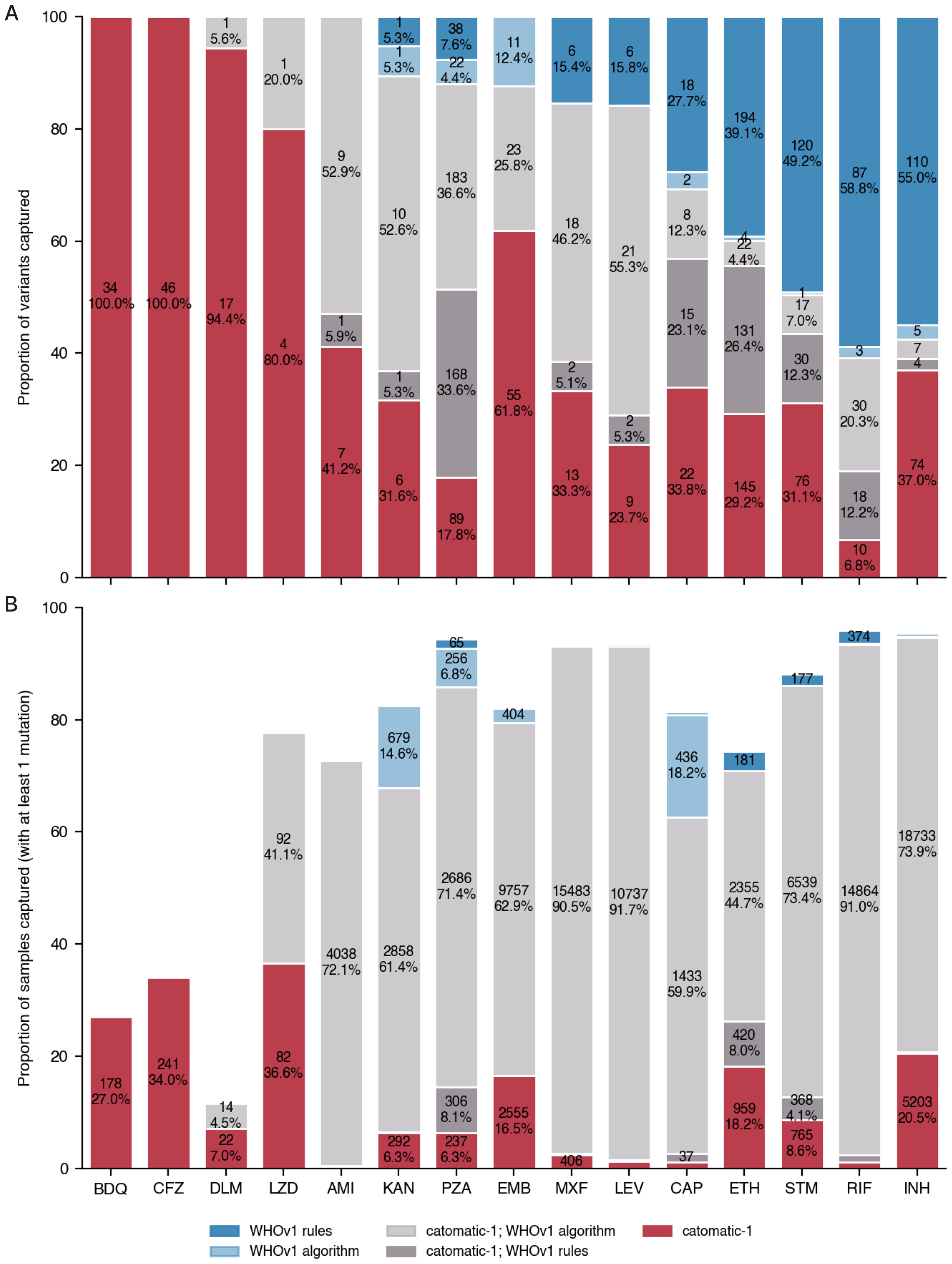


Figure S7: Proportion of the Training Set captured by `WHOv1`'s algorithm, its rules, and `catomatic-1`, with mutations or samples in both coloured grey. A. Bars show the proportion of mutations classified by `WHOv1` and `catomatic-1`, stratifying by what the `WHO`'s algorithm captured (light hues) and what their Expert Rules added (dark hues), ordered by the total proportions achieved by `catomatic-1` (i.e the top the of the light grey bars). B. The corresponding proportions of samples in the Training Set (and thus all having at least 1 mutation in relevant genes) for which predictions can be made by the same groups in A - i.e their realised effect. Pairwise classification summaries are in Table S6.

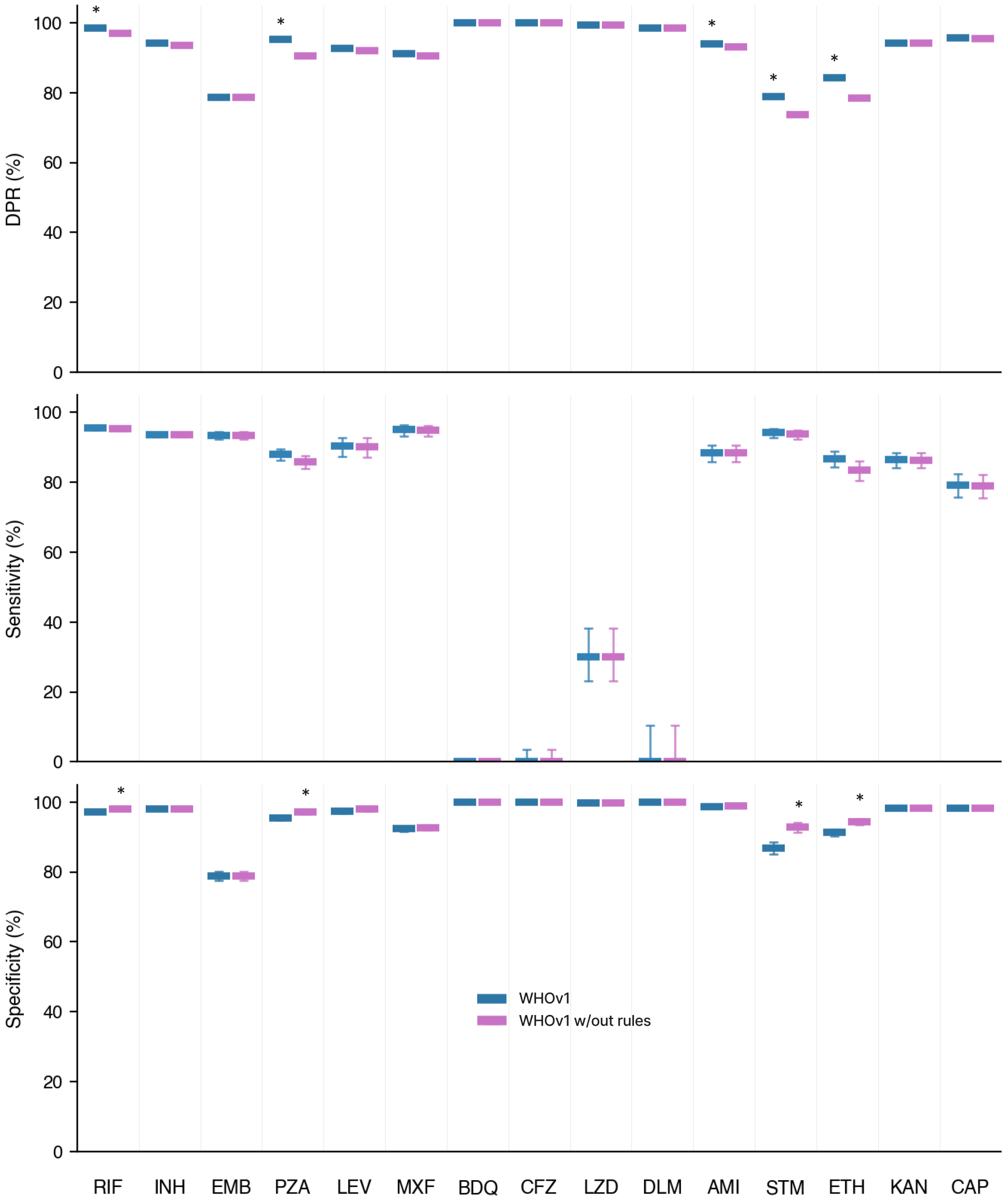


Figure S8: Sensitivity, specificity, and definitive prediction rate (DPR) for `WHOv1` (blue) and `WHOv1` with all rules removed and additional grading classifications reverted (violet), evaluated on The Validation Dataset of 14,380 samples. The significantly larger value at the 95% confidence level is asterisked. Point values are in Table S4.

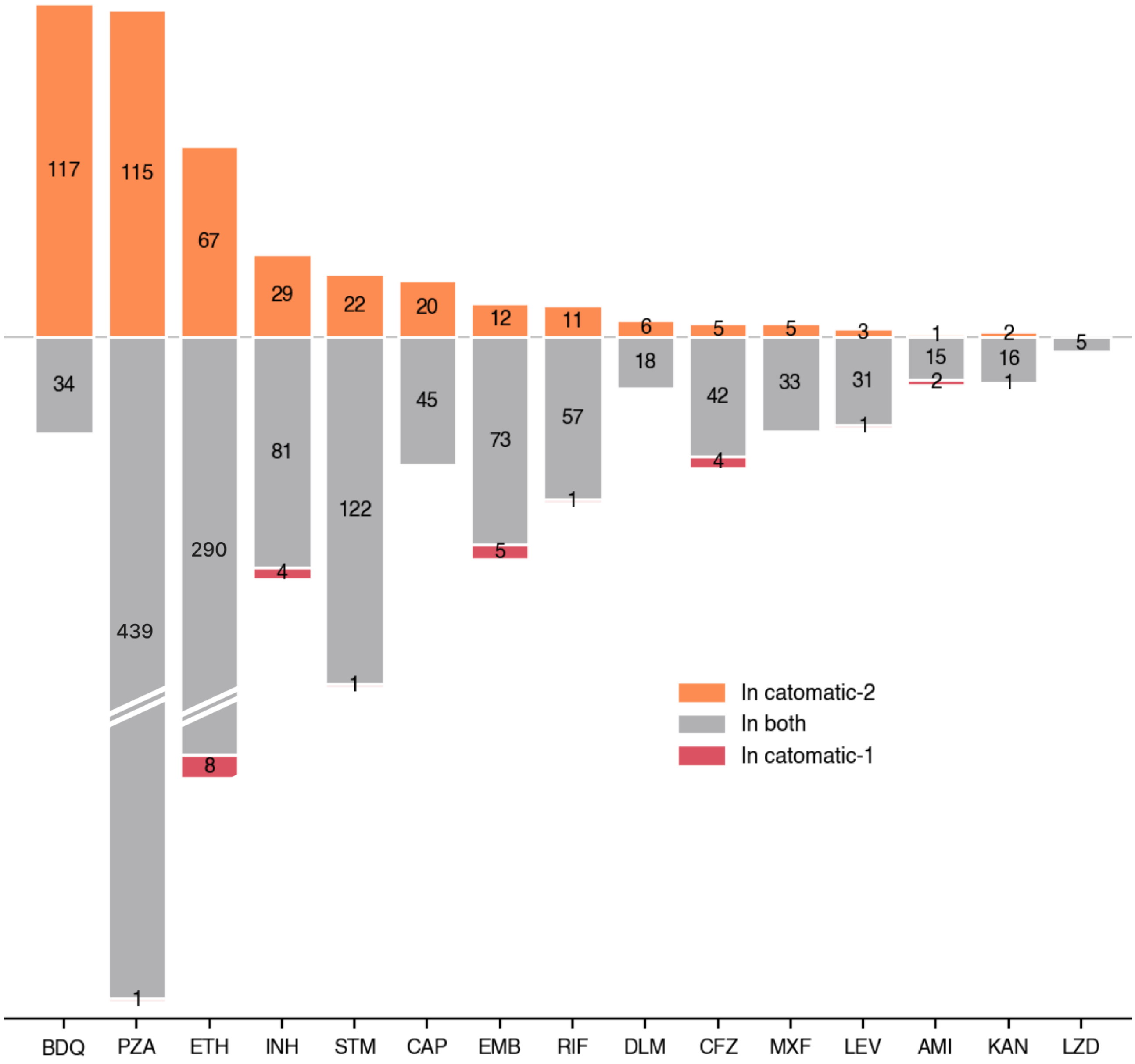


Figure S9: Bars show the number of unique mutations in The Training Dataset assigned differently across `catomatic-1` and `catomatic-2`. Mutations present only in `catomatic-1` are shown in red. Mutations present in both catalogues are shown in grey. Mutations present only in `catomatic-2` are shown in orange. Bars below the axis for PZA and ETH are truncated due to their size. The per-variant classifications for catomatic-1 and catomatic-2 are compared in the supplementary spreadsheet

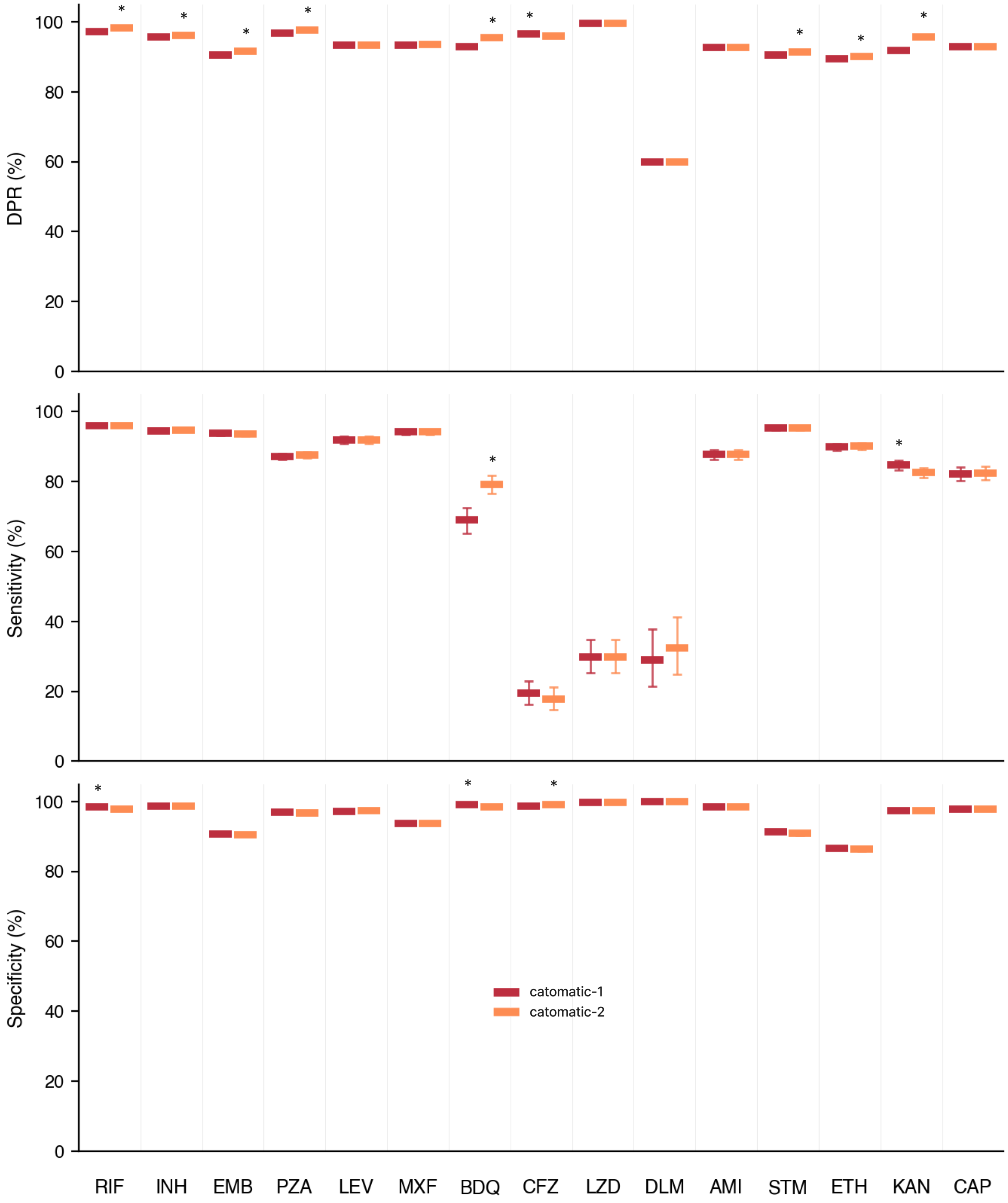


Figure S10: Sensitivity, specificity, and definitive prediction rate (DPR) for `catomatic-1` (red) and `catomatic-2` (orange) evaluated on the Entire Dataset (Training + Validation). The significantly larger value at the 95% confidence level is asterisked. Note this not an independent measure of performance for either catalogue, as `catomatic-2` was built on The Entire Dataset and `catomatic-1` was built on a subset. Plotted values correspond to those in Table S7.

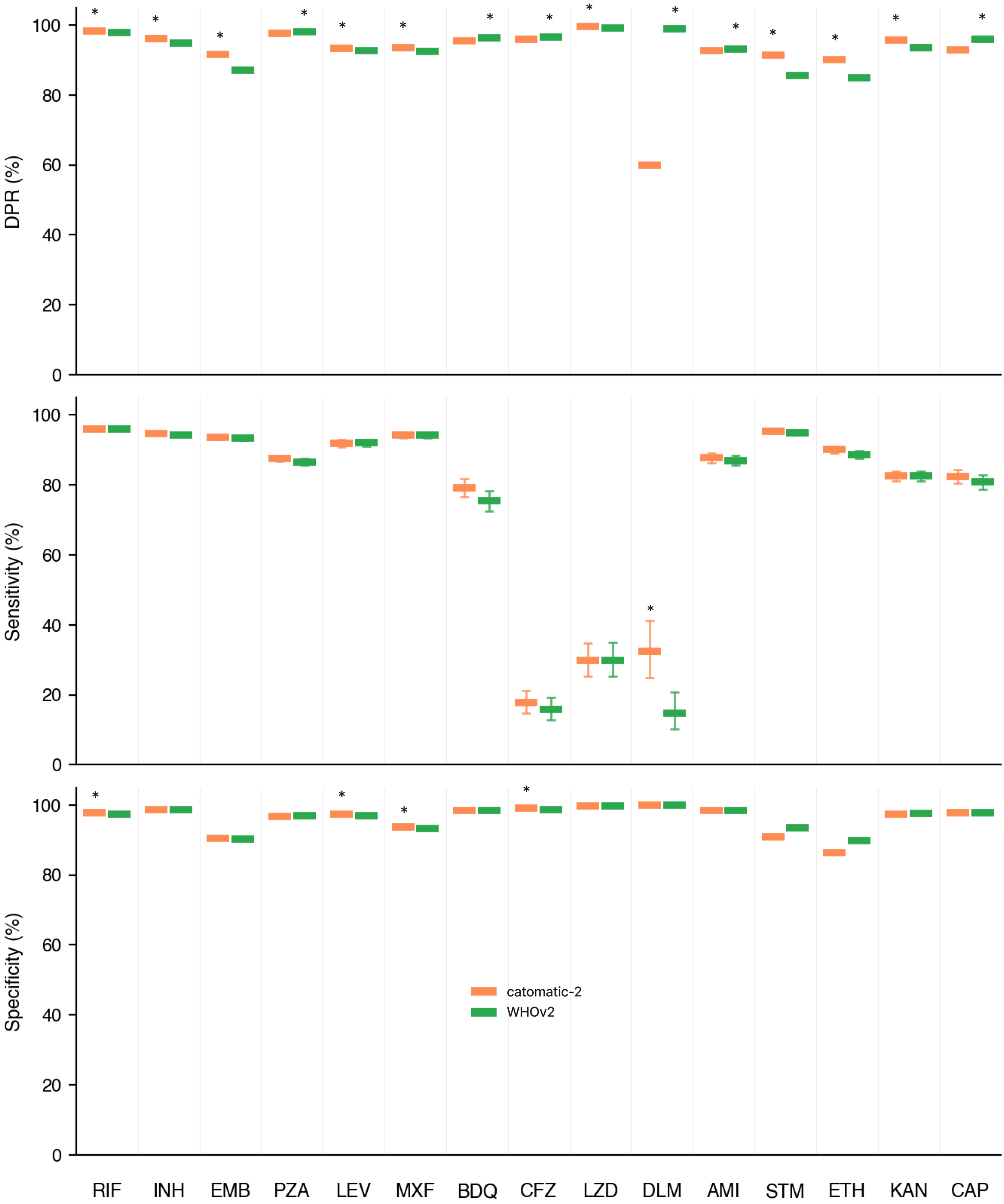


Figure S11: Sensitivity, specificity, and definitive prediction rate (DPR) for `catomatic-2` (orange) and `WHOv2` (green) evaluated on the Entire Dataset (Training + Validation). The significantly larger value at the 95% confidence level is asterisked. Note this not an independent measure of performance for either catalogue, as `catomatic-2` was built on The Entire Dataset and `WHOv2` was likely built on many of these same samples. Plotted values correspond to those in Table S7.

| Drug | Comparison | Sensitivity | | Specificity | | DPR | |
|---|---|---|---|---|---|---|---|
| | | Δ (%) | p | Δ (%) | p | Δ (%) | p |
| RIF | Build: 0.1 vs 0.9 | 0.00 | 1.000 | 0.00 | 0.974 | 0.16 | 0.194 |
| | Test: 0.1 vs 0.9 | **+2.12** | **0.001** | -0.13 | 0.212 | -0.22 | 0.074 |
| INH | Build: 0.1 vs 0.9 | -0.11 | 0.682 | 0.06 | 0.572 | **+1.92** | **0.001** |
| | Test: 0.1 vs 0.9 | **+1.71** | **0.001** | -0.14 | 0.167 | **-0.31** | **0.034** |
| EMB | Build: 0.1 vs 0.9 | -0.66 | 0.121 | **+0.58** | **0.019** | **+4.76** | **0.001** |
| | Test: 0.1 vs 0.9 | **+2.81** | **0.001** | **-0.50** | **0.032** | **-1.02** | **0.001** |
| PZA | Build: 0.1 vs 0.9 | 0.00 | 0.932 | 0.00 | 0.850 | 0.00 | 0.474 |
| | Test: 0.1 vs 0.9 | **+5.45** | **0.001** | **-0.36** | **0.032** | -0.26 | 0.065 |
| LEV | Build: 0.1 vs 0.9 | -0.00 | 0.996 | 0.00 | 1.000 | -0.01 | 0.979 |
| | Test: 0.1 vs 0.9 | **+8.29** | **0.001** | -0.36 | 0.135 | -0.54 | 0.095 |
| MXF | Build: 0.1 vs 0.9 | 0.00 | 1.000 | 0.00 | 1.000 | 0.00 | 1.000 |
| | Test: 0.1 vs 0.9 | **+7.06** | **0.001** | **-0.63** | **0.027** | **-0.55** | **0.038** |
| BDQ | Build: 0.1 vs 0.9 | 6.37 | 0.309 | **-0.38** | **0.001** | 0.51 | 0.066 |
| | Test: 0.1 vs 0.9 | **+20.59** | **0.001** | **-0.35** | **0.004** | **-0.55** | **0.034** |
| CFZ | Build: 0.1 vs 0.9 | **+6.15** | **0.016** | **-0.85** | **0.001** | **+1.04** | **0.001** |
| | Test: 0.1 vs 0.9 | **+7.41** | **0.003** | **-0.40** | **0.003** | **-0.50** | **0.026** |
| LZD | Build: 0.1 vs 0.9 | 0.00 | 1.000 | 0.00 | 1.000 | 0.00 | 1.000 |
| | Test: 0.1 vs 0.9 | +3.90 | 0.378 | -0.06 | 0.266 | -0.03 | 0.604 |
| DLM | Build: 0.1 vs 0.9 | 1.48 | 0.833 | 0.00 | 1.000 | 0.02 | 0.977 |
| | Test: 0.1 vs 0.9 | +2.20 | 0.752 | +0.72 | 0.720 | -1.01 | 0.131 |
| AMI | Build: 0.1 vs 0.9 | 0.00 | 1.000 | 0.00 | 1.000 | 0.00 | 1.000 |
| | Test: 0.1 vs 0.9 | **+5.91** | **0.001** | -0.22 | 0.109 | -0.46 | 0.091 |
| STM | Build: 0.1 vs 0.9 | 0.01 | 0.985 | -0.06 | 0.868 | 0.10 | 0.768 |
| | Test: 0.1 vs 0.9 | **+2.42** | **0.001** | -0.27 | 0.479 | -0.35 | 0.287 |
| ETH | Build: 0.1 vs 0.9 | 0.00 | 1.000 | 0.37 | 0.434 | 0.00 | 0.394 |
| | Test: 0.1 vs 0.9 | **+3.87** | **0.001** | -0.84 | 0.069 | **-0.81** | **0.021** |
| KAN | Build: 0.1 vs 0.9 | 0.02 | 0.988 | -0.01 | 0.946 | 0.02 | 0.939 |
| | Test: 0.1 vs 0.9 | **+5.24** | **0.001** | -0.22 | 0.243 | -0.42 | 0.141 |
| CAP | Build: 0.1 vs 0.9 | -0.25 | 0.885 | 0.06 | 0.787 | **+2.16** | **0.001** |
| | Test: 0.1 vs 0.9 | **+7.75** | **0.001** | **-0.44** | **0.035** | -0.42 | 0.220 |

Table S3: Differences in sensitivity, specificity, and definitive prediction rate (DPR) when applying an $FRS_{min}$ of 0.1 rather than 0.9 at catalogue build time or when making predictions (the corresponding test and build $FRS_{min}$, respectively, are held at 0.1), on The Training Dataset. Significant results ($p < 0.05$) are shown in bold. p-values below 0.001 are rounded to 0.001.

| | WHOv1 | | | WHOv1 w/out rules | | |
|---|---|---|---|---|---|---|
| Drug | Sens | Spec | DPR | Sens | Spec | DPR |
| RIF | $95.5^{+0.58}_{-0.66}$ | $97.1^{+0.36}_{-0.42}$ | $98.6^{+0.20}_{-0.23}$ | $95.4^{+0.59}_{-0.68}$ | $97.9^{+0.31}_{-0.36}$ | $96.9^{+0.30}_{-0.33}$ |
| INH | $93.7^{+0.64}_{-0.71}$ | $97.9^{+0.33}_{-0.40}$ | $94.1^{+0.42}_{-0.45}$ | $93.6^{+0.65}_{-0.72}$ | $98.0^{+0.33}_{-0.39}$ | $93.5^{+0.44}_{-0.47}$ |
| EMB | $93.5^{+1.04}_{-1.22}$ | $78.8^{+1.29}_{-1.35}$ | $78.7^{+0.94}_{-0.97}$ | $93.5^{+1.04}_{-1.22}$ | $78.8^{+1.29}_{-1.35}$ | $78.7^{+0.94}_{-0.97}$ |
| PZA | $88.0^{+1.49}_{-1.68}$ | $95.4^{+0.54}_{-0.61}$ | $95.2^{+0.47}_{-0.53}$ | $85.9^{+1.73}_{-1.93}$ | $97.1^{+0.43}_{-0.51}$ | $90.6^{+0.66}_{-0.71}$ |
| LEV | $90.4^{+2.41}_{-3.11}$ | $97.4^{+0.53}_{-0.67}$ | $92.7^{+0.83}_{-0.91}$ | $90.3^{+2.45}_{-3.17}$ | $98.0^{+0.46}_{-0.60}$ | $92.0^{+0.86}_{-0.96}$ |
| MXF | $95.0^{+1.32}_{-1.76}$ | $92.3^{+0.64}_{-0.71}$ | $91.0^{+0.63}_{-0.67}$ | $94.9^{+1.33}_{-1.79}$ | $92.7^{+0.63}_{-0.69}$ | $90.5^{+0.64}_{-0.69}$ |
| BDQ | $0^{+0.46}_{-0}$ | $100^{+0}_{-0.11}$ | $100^{+0}_{-0.09}$ | $0^{+0.46}_{-0}$ | $100^{+0}_{-0.11}$ | $100^{+0}_{-0.09}$ |
| CFZ | $0^{+3.50}_{-0}$ | $100^{+0}_{-0.11}$ | $100^{+0}_{-0.11}$ | $0^{+3.50}_{-0}$ | $100^{+0}_{-0.11}$ | $100^{+0}_{-0.11}$ |
| LZD | $30.1^{+8.17}_{-7.08}$ | $99.7^{+0.12}_{-0.22}$ | $99.3^{+0.21}_{-0.29}$ | $30.1^{+8.17}_{-7.08}$ | $99.7^{+0.12}_{-0.22}$ | $99.3^{+0.21}_{-0.29}$ |
| DLM | $0^{+10.4}_{-0}$ | $100^{+0}_{-0.14}$ | $98.4^{+0.40}_{-0.53}$ | $0^{+10.4}_{-0}$ | $100^{+0}_{-0.14}$ | $98.4^{+0.40}_{-0.53}$ |
| AMI | $88.5^{+2.12}_{-2.53}$ | $98.8^{+0.24}_{-0.30}$ | $93.9^{+0.51}_{-0.55}$ | $88.4^{+2.13}_{-2.53}$ | $98.8^{+0.24}_{-0.29}$ | $93.0^{+0.55}_{-0.58}$ |
| STM | $94.2^{+1.14}_{-1.39}$ | $86.8^{+1.61}_{-1.81}$ | $78.9^{+1.30}_{-1.37}$ | $93.8^{+1.22}_{-1.49}$ | $92.8^{+1.24}_{-1.47}$ | $73.7^{+1.42}_{-1.46}$ |
| ETH | $86.7^{+2.09}_{-2.40}$ | $91.2^{+0.93}_{-1.02}$ | $84.2^{+1.00}_{-1.05}$ | $83.5^{+2.56}_{-2.91}$ | $94.5^{+0.75}_{-0.86}$ | $78.4^{+1.14}_{-1.18}$ |
| KAN | $86.4^{+2.01}_{-2.29}$ | $98.2^{+0.29}_{-0.35}$ | $94.2^{+0.49}_{-0.53}$ | $86.3^{+2.02}_{-2.30}$ | $98.2^{+0.29}_{-0.35}$ | $94.1^{+0.49}_{-0.54}$ |
| CAP | $79.3^{+3.18}_{-3.58}$ | $98.1^{+0.33}_{-0.40}$ | $95.6^{+0.49}_{-0.55}$ | $79.0^{+3.21}_{-3.61}$ | $98.2^{+0.32}_{-0.40}$ | $95.4^{+0.50}_{-0.55}$ |

Table S4: Sensitivity (Sens) (%), specificity (Spec) (%), and definite prediction rate (DPR) (%) for `WHOv1` (with rules) and `WHOv1` without rules across drugs using the ternary prediction system, applied to The Validation Dataset. Values are plotted in Fig S8 and are shown here as point estimates with confidence intervals, computed from the reported lower and upper confidence limits as $x^{+\text{upper}-x}_{-x+\text{lower}}$. Differences in performance and p-values are in Table S5. PPV values and upper/lower bounds are omitted for clarity but are available in the expanded Table S4 in the supplementary spreadsheet.

| Drug | Comparison | Sensitivity | | Specificity | | DPR | |
|---|---|---|---|---|---|---|---|
| | | Δ (%) | p | Δ (%) | p | Δ (%) | p |
| RIF | catomatic-1 - WHOv1 no rules | +0.04 | 0.929 | -0.05 | 0.832 | **+0.69** | **0.002** |
| | catomatic-1 - WHOv1 | -0.10 | 0.820 | **+0.78** | **0.003** | **-0.99** | **0.001** |
| | WHOv1 - WHOv1 no rules | +0.14 | 0.753 | **-0.83** | **0.002** | **+1.68** | **0.001** |
| INH | catomatic-1 - WHOv1 no rules | **+1.01** | **0.032** | -0.33 | 0.225 | **+2.21** | **0.001** |
| | catomatic-1 - WHOv1 | **+0.93** | **0.047** | -0.24 | 0.375 | **+1.63** | **0.001** |
| | WHOv1 - WHOv1 no rules | +0.08 | 0.873 | -0.09 | 0.745 | +0.58 | 0.070 |
| EMB | catomatic-1 - WHOv1 no rules | **-2.00** | **0.021** | **+3.56** | **0.001** | **+10.0** | **0.001** |
| | catomatic-1 - WHOv1 | **-2.00** | **0.021** | **+3.56** | **0.001** | **+10.0** | **0.001** |
| | WHOv1 - WHOv1 no rules | 0.00 | 1.000 | 0.00 | 1.000 | 0.00 | 1.000 |
| PZA | catomatic-1 - WHOv1 no rules | **+2.70** | **0.026** | **-1.05** | **0.004** | **+3.53** | **0.001** |
| | catomatic-1 - WHOv1 | +0.66 | 0.561 | +0.65 | 0.108 | **-1.08** | **0.005** |
| | WHOv1 - WHOv1 no rules | +2.04 | 0.096 | **-1.69** | **0.001** | **+4.60** | **0.001** |
| LEV | catomatic-1 - WHOv1 no rules | -0.07 | 0.971 | -0.27 | 0.487 | +0.93 | 0.144 |
| | catomatic-1 - WHOv1 | -0.25 | 0.900 | +0.33 | 0.424 | +0.20 | 0.744 |
| | WHOv1 - WHOv1 no rules | +0.18 | 0.929 | -0.61 | 0.136 | +0.73 | 0.257 |
| MXF | catomatic-1 - WHOv1 no rules | -0.16 | 0.883 | +0.06 | 0.899 | **+2.73** | **0.001** |
| | catomatic-1 - WHOv1 | -0.25 | 0.823 | +0.44 | 0.352 | **+2.16** | **0.001** |
| | WHOv1 - WHOv1 no rules | +0.08 | 0.940 | -0.38 | 0.425 | +0.57 | 0.234 |
| BDQ | catomatic-1 - WHOv1 no rules | **+73.6** | **0.001** | **-1.16** | **0.001** | **-14.5** | **0.001** |
| | catomatic-1 - WHOv1 | **+73.6** | **0.001** | **-1.16** | **0.001** | **-14.5** | **0.001** |
| | WHOv1 - WHOv1 no rules | 0.00 | NaN | 0.00 | NaN | 0.00 | NaN |
| CFZ | catomatic-1 - WHOv1 no rules | **+19.2** | **0.001** | **-1.28** | **0.001** | **-3.35** | **0.001** |
| | catomatic-1 - WHOv1 | **+19.2** | **0.001** | **-1.28** | **0.001** | **-3.35** | **0.001** |
| | WHOv1 - WHOv1 no rules | 0.00 | NaN | 0.00 | NaN | 0.00 | NaN |
| LZD | catomatic-1 - WHOv1 no rules | -0.22 | 0.968 | 0.00 | 0.995 | +0.27 | 0.095 |
| | catomatic-1 - WHOv1 | -0.22 | 0.968 | 0.00 | 0.995 | +0.27 | 0.095 |
| | WHOv1 - WHOv1 no rules | 0.00 | 1.000 | 0.00 | 1.000 | 0.00 | 1.000 |
| DLM | catomatic-1 - WHOv1 no rules | +8.70 | 0.085 | 0.00 | NaN | **-44.5** | **0.001** |
| | catomatic-1 - WHOv1 | +8.70 | 0.085 | 0.00 | NaN | **-44.5** | **0.001** |
| | WHOv1 - WHOv1 no rules | 0.00 | NaN | 0.00 | NaN | 0.00 | 1.000 |
| AMI | catomatic-1 - WHOv1 no rules | +0.02 | 0.992 | -0.06 | 0.743 | **+0.88** | **0.026** |
| | catomatic-1 - WHOv1 | 0.00 | 1.000 | -0.03 | 0.877 | +0.03 | 0.947 |
| | WHOv1 - WHOv1 no rules | +0.02 | 0.992 | -0.03 | 0.862 | **+0.86** | **0.030** |
| STM | catomatic-1 - WHOv1 no rules | +1.01 | 0.256 | **-11.1** | **0.001** | **+12.4** | **0.001** |
| | catomatic-1 - WHOv1 | +0.60 | 0.485 | **-5.08** | **0.001** | **+7.25** | **0.001** |
| | WHOv1 - WHOv1 no rules | +0.41 | 0.663 | **-5.99** | **0.001** | **+5.13** | **0.001** |
| ETH | catomatic-1 - WHOv1 no rules | +3.31 | 0.064 | **-8.71** | **0.001** | **+10.1** | **0.001** |
| | catomatic-1 - WHOv1 | +0.08 | 0.963 | **-5.50** | **0.001** | **+4.34** | **0.001** |
| | WHOv1 - WHOv1 no rules | +3.23 | 0.071 | **-3.21** | **0.001** | **+5.78** | **0.001** |
| KAN | catomatic-1 - WHOv1 no rules | +0.43 | 0.784 | -0.46 | 0.061 | **-2.48** | **0.001** |
| | catomatic-1 - WHOv1 | +0.36 | 0.818 | -0.46 | 0.061 | **-2.54** | **0.001** |
| | WHOv1 - WHOv1 no rules | +0.07 | 0.964 | 0.00 | 1.000 | +0.06 | 0.866 |
| CAP | catomatic-1 - WHOv1 no rules | +0.30 | 0.901 | -0.09 | 0.722 | **-1.55** | **0.001** |
| | catomatic-1 - WHOv1 | +0.08 | 0.975 | -0.04 | 0.886 | **-1.70** | **0.001** |
| | WHOv1 - WHOv1 no rules | +0.23 | 0.926 | -0.06 | 0.831 | +0.15 | 0.693 |

Table S5: Performance differences between `catomatic-1` and `WHOv1` and `WHOv1 no rules`, calculated on The Validation Set and corresponding to values in Table 3 and Table S4. Significant results ($p < 0.05$) are shown in bold. p-values less than 0.001 are rounded to 0.001.

| | catomatic: WHOv1 | | | | | | | | | catomatic: WHOv1 algorithm | | | | | | | | |
|---|---|---|---|---|---|---|---|---|---|---|---|---|---|---|---|---|---|---|
| **Drug** | **R.R** | **R.S** | **R.U** | **S.R** | **S.S** | **S.U** | **U.R** | **U.S** | **Total** | **R.R** | **R.S** | **R.U** | **S.R** | **S.S** | **S.U** | **U.R** | **U.S** | **Total** |
| AMI | 4 | 0 | 7 | 0 | 6 | 0 | 0 | 0 | 17 | 3 | 0 | 8 | 0 | 6 | 0 | 0 | 0 | 17 |
| BDQ | 0 | 0 | 34 | 0 | 0 | 0 | 0 | 0 | 34 | 0 | 0 | 34 | 0 | 0 | 0 | 0 | 0 | 34 |
| CAP | 21 | 0 | 22 | 0 | 2 | 0 | 18 | 2 | 65 | 7 | 0 | 36 | 0 | 2 | 0 | 0 | 2 | 47 |
| CFZ | 0 | 0 | 46 | 0 | 0 | 0 | 0 | 0 | 46 | 0 | 0 | 46 | 0 | 0 | 0 | 0 | 0 | 46 |
| DLM | 1 | 0 | 17 | 0 | 0 | 0 | 0 | 0 | 18 | 1 | 0 | 17 | 0 | 0 | 0 | 0 | 0 | 18 |
| EMB | 12 | 0 | 8 | 0 | 11 | 47 | 3 | 8 | 89 | 12 | 0 | 8 | 0 | 11 | 47 | 3 | 8 | 89 |
| ETH | 153 | 0 | 145 | 0 | 0 | 0 | 198 | 0 | 496 | 61 | 0 | 237 | 0 | 0 | 0 | 4 | 0 | 302 |
| INH | 10 | 0 | 52 | 0 | 1 | 22 | 112 | 3 | 200 | 6 | 0 | 56 | 0 | 1 | 22 | 2 | 3 | 90 |
| KAN | 8 | 0 | 3 | 0 | 3 | 3 | 0 | 2 | 19 | 7 | 0 | 4 | 0 | 3 | 3 | 0 | 1 | 18 |
| LEV | 10 | 0 | 3 | 0 | 13 | 6 | 4 | 2 | 38 | 8 | 0 | 5 | 0 | 13 | 6 | 0 | 0 | 32 |
| LZD | 1 | 0 | 0 | 0 | 0 | 4 | 0 | 0 | 5 | 1 | 0 | 0 | 0 | 0 | 4 | 0 | 0 | 5 |
| MXF | 10 | 0 | 1 | 0 | 10 | 12 | 4 | 2 | 39 | 8 | 0 | 3 | 0 | 10 | 12 | 0 | 0 | 33 |
| PZA | 350 | 1 | 89 | 0 | 0 | 0 | 46 | 14 | 500 | 252 | 1 | 187 | 0 | 0 | 0 | 5 | 14 | 462 |
| RIF | 40 | 0 | 2 | 1 | 7 | 8 | 88 | 2 | 148 | 23 | 0 | 19 | 1 | 7 | 8 | 1 | 2 | 61 |
| STM | 43 | 0 | 72 | 0 | 4 | 4 | 121 | 0 | 244 | 15 | 0 | 100 | 0 | 4 | 4 | 1 | 0 | 124 |

Table S6: Pairwise comparison of mutation classifications between `catomatic-1` and `WHOv1`, showing the number of mutations assigned to each pair of classifications (e.g. R.R indicates variants classified as resistance-associated (R) by both catalogues; R.S variants classified as resistance-associated by `catomatic` and not-associated (S) by `WHOv1`; U.R variants unclassified by `catomatic` but classified as resistance-associated by `WHOv1`, etc.). The left panel compares `catomatic` to `WHOv1` including expert rules, while the right panel compares `catomatic` to the `WHOv1` algorithm excluding expert rules. Mutation-level pairwise differences are available in the Supplementary Spreadsheet

| | catomatic-1 | | | catomatic-2 | | | WHOv1 | | | WHOv2 | | |
|---|---|---|---|---|---|---|---|---|---|---|---|---|
| Drug | Sens | Spec | DPR | Sens | Spec | DPR | Sens | Spec | DPR | Sens | Spec | DPR |
| RIF | 95.95 | 98.39 | 97.22 | 96.01 | 97.72 | 98.21 | 96.02 | 97.42 | 97.96 | 96.05 | 97.41 | 97.77 |
| INH | 94.56 | 98.56 | 95.70 | 94.60 | 98.55 | 96.07 | 93.52 | 98.72 | 95.34 | 94.22 | 98.61 | 94.76 |
| EMB | 93.85 | 90.61 | 90.47 | 93.63 | 90.46 | 91.57 | 95.15 | 89.28 | 83.74 | 93.43 | 90.22 | 87.15 |
| PZA | 87.15 | 97.00 | 96.75 | 87.56 | 96.68 | 97.56 | 86.32 | 96.80 | 97.41 | 86.55 | 96.96 | 97.97 |
| LEV | 91.96 | 97.26 | 93.19 | 91.94 | 97.35 | 93.38 | 92.02 | 97.00 | 92.46 | 92.07 | 96.85 | 92.73 |
| MXF | 94.22 | 93.68 | 93.19 | 94.22 | 93.71 | 93.54 | 94.30 | 93.23 | 91.13 | 94.28 | 93.19 | 92.40 |
| BDQ | 68.96 | 98.99 | 92.92 | 79.25 | 98.46 | 95.39 | 0.00 | 100.00 | 100.00 | 75.42 | 98.42 | 96.29 |
| CFZ | 19.40 | 98.70 | 96.44 | 17.71 | 99.13 | 95.97 | 0.00 | 100.00 | 100.00 | 15.79 | 98.57 | 96.44 |
| LZD | 29.82 | 99.77 | 99.64 | 29.82 | 99.77 | 99.64 | 29.91 | 99.77 | 99.14 | 29.91 | 99.77 | 99.14 |
| DLM | 28.95 | 99.94 | 59.93 | 32.50 | 99.90 | 60.00 | 6.83 | 100.00 | 98.78 | 14.77 | 99.95 | 98.92 |
| AMI | 87.78 | 98.41 | 92.62 | 87.77 | 98.42 | 92.60 | 87.74 | 98.43 | 92.57 | 87.02 | 98.44 | 93.06 |
| STM | 95.26 | 91.43 | 90.41 | 95.35 | 90.92 | 91.44 | 94.84 | 93.36 | 85.72 | 94.82 | 93.52 | 85.53 |
| ETH | 89.86 | 86.68 | 89.33 | 90.14 | 86.40 | 90.08 | 89.14 | 90.13 | 85.40 | 88.75 | 89.91 | 85.00 |
| KAN | 84.69 | 97.31 | 91.77 | 82.64 | 97.44 | 95.59 | 82.49 | 97.69 | 94.03 | 82.52 | 97.67 | 93.41 |
| CAP | 82.27 | 97.77 | 92.84 | 82.49 | 97.76 | 92.96 | 80.39 | 97.79 | 95.82 | 80.88 | 97.74 | 95.83 |

Table S7: Sensitivity (Sens) (%), specificity (Spec) (%), and definite prediction rate (DPR) (%) point values of `catomatic-1`, `catomatic-2`, `WHOv1`, and `WHOv2` across drugs using the ternary prediction system, applied to the Entire Dataset (Training + Validation). All catalogues have seen samples in The Entire Dataset, therefore this is not an independent measure of performance. `catomatic-2` and `WHOv2` were trained on more samples (in The Validation Dataset) than `catomatic-1` and `WHOv1`. PPV values, and confidence limits for all metrics are omitted for clarity but are available in the Supplementary Material Spreadsheet.
